# KF4 anti-CELA1 Antibody and Purified α1-Antitrypsin Have Similar but Not Additive Efficacy in Preventing Emphysema in Murine α1-Antitrypsin Deficiency

**DOI:** 10.1101/2024.05.07.592994

**Authors:** Andrew J. Devine, Noah J. Smith, Rashika Joshi, Brandon Brooks-Patton, Jenna Dunham, Ansley N. Varisco, Emily M. Goodman, Qiang Fan, Basilia Zingarelli, Brian M. Varisco

**Affiliations:** Ohio University Heritage College of Osteopathic Medicine, Athens, OH, USA; College of Medicine, University of Cincinnati, Cincinnati, OH, USA; Critical Care Medicine, Cincinnati Children’s Hospital Medical Center, Cincinnati, OH, USA; Northern Kentucky University, Covington, KY, USA; University of Arkansas at Little Rock, Little Rock, AR, USA; University of Arkansas for Medical Sciences, Little Rock, AR, USA; Arkansas Children’s Research Institute, Little Rock, AR, USA

**Keywords:** Protease, anti-Protease, Elastase, Emphysema, COPD, Alpha-1 Antitrypsin

## Abstract

Alpha-1 antitrypsin (AAT) deficiency is the most common genetic cause of emphysema. Chymotrypsin-like Elastase 1 (CELA1) is a serine protease neutralized by AAT and is important in emphysema progression. *Cela1*-deficiency is protective in a murine models of AAT-deficient emphysema. KF4 anti-CELA1 antibody prevented emphysema in PPE and cigarette smoke models in wild type mice. We evaluated potential toxicities of KF4 and its ability to prevent emphysema in AAT deficiency. We found Cela1 protein expression in mouse lung, pancreas, small intestine, and spleen. In toxicity studies, mice treated with KF4 25 mg/kg weekly for four weeks showed an elevation in blood urea nitrogen and slower weight gain compared to lower doses or equivalent dose IgG. In histologic grading of tissue injury of the lung, kidney, liver, and heart, there was some evidence of liver injury with KF4 25 mg/kg, but in all tissues, injury was less than in control mice subjected to cecal ligation and puncture. In efficacy studies, KF4 doses as low as 0.5 mg/kg reduced the lung elastase activity of *AAT^-/-^*mice treated with 0.2 units of PPE. In this injury model, *AAT^-/-^*mice treated with KF4 1 mg/kg weekly, human purified AAT 60 mg/kg weekly, and combined KF4 and AAT treatment had less emphysema than mice treated with IgG 1 mg/kg weekly. However, the efficacy of KF4, AAT, or KF4 & AAT was similar. While KF4 might be an alternative to AAT replacement, combined KF4 and AAT replacement does not confer additional benefit.

## Introduction

Alpha-1 antitrypsin (AAT) deficiency is the most common genetic factor predisposing people to emphysema (1). Patients typically present in the fourth or fifth decade of life, and despite meta-analyses suggesting only marginal benefit (2), replacement therapy with purified, human AAT is standard of care (3).

AAT is an anti-protease that neutralizes Neutrophil Elastase, Cathepsin G, and Proteinase-3 (4, 5). Our group has shown that AAT also neutralizes and an alveolar type 2 cell-secreted serine protease called Chymotrypsin-like Elastase 1 (CELA1) (6–8). CELA1 plays a physiologic role in the postnatal lung by reducing lung elastase (6) and at baseline *Cela1^-/-^* mice have slightly smaller alveoli than wild type mice. In antisense oligonucleotide (6) and genetic (7) models of AAT-deficiency, *Cela1^-/-^* AAT-deficient mice are protected from emphysema. CELA1-AAT protein complexes can be found in human lung indicating that AAT likely neutralized CELA1 by covalent binding (8).

The KF4 anti-CELA1 monoclonal antibody is a mouse IgG1 that covalently binds to a region of the CELA1 molecule containing the catalytic triad histidine (8). KF4 protects wild type mice from cigarette-smoke induced emphysema and from emphysema progression following tracheal administration of porcine pancreatic elastase (PPE). Based upon the premise that AAT neutralization of CELA1 plays a key role in the pathogenesis of emphysema in AAT-deficiency, we sought to test whether Cela1-neutralization using KF4 antibody could protect AAT-deficient mice from emphysema and compare its efficacy with AAT replacement therapy.

## Methods

### Animal Use

Animal use was approved by the Cincinnati Children’s Hospital Institutional Review board under authorization 2020-0054 (BMV). In addition, we utilized previously collected images from mice utilized under authorization IACUC2021-0087 (BZ). Results from these studies were previously published (9, 10).

### Tracheal PPE Administration

Eight to ten week-old male and female C57BL6 mice were anesthetized with 2% isoflurane and suspended by the incisors on a mouse intubating board. The tongue was withdrawn using forceps, and 0.2 units of PPE (Sigma Aldrich, E1250), in 100 µL of PBS was instilled into the oropharyngeal cavity. The nose was occluded with a finger and the mouse remained suspended until began recovering and aspirated the solution. The mouse was recovered in a cage and observed until ambulation resumed.

### Mouse Cecal Ligation and Puncture

As described previously (9) mice were anesthetized with isoflurane and the cecum visualized through a midline incision The cecum was ligated with 3-0 silk and a single through-and-through puncture made with a 22 gauge needle. Cecal contents were expressed and the abdomen was closed with prolene suture. For sham surgeries, the intestines were manipulated without ligation or puncture. The mice were treated with buprenorphine and recovered. At 6 and 18 hours, mice were sacrificed and vital organs collected.

### Tissue Processing

All tissues were fixed in 4% paraformaldehyde in PBS overnight, brought into 100% ethanol by serial dehydration and paraffinized. For liver specimens, only the left lobe was placed in the cassette for embedding. Only the left kidney was embedded. Whole hearts were embedded but only left ventricles assessed. For lung lobes, the four right lung lobes were arranged in a cassette with random orientation and embedded. Five micron sections of each specimen were mounted on a slide and stained using hematoxylin and eosin.

### Imaging

Using a Nikon Ti2 microscope, for liver, kidney, and heart tissues, five 10X and five 40X images per slide were obtained. For lung images, five 10X images from each lobe were obtained and a total of five 40X images from each lung lobe were obtained.

### Mean Linear Intercept Analysis

Using the methods of Dunnill et al (11), after de-identification, mean linear intercept values were determined for each 10X lung photomicrograph by a reviewer (AD).

### Immunohistochemistry

For immunohistochemistry the ABC Vectastain kit was used with a previously validated anti-CELA1 guinea pig polyclonal antibody (5).

### Toxicity Assessment

Procedure: Eight week-old wild type female C57BL6 mice were administered 5, 12.5, or 25 mg/kg KF4 or 25 mg/kg mouse anti-Human IgG (Jackson Laboratories, 711-005-152) peritoneally. For 5, 12.5, and 25 mg/kg doses, a maximum of 5 mg/kg was administered at any given time and antibody was administered 1, 3, and 5 times per week respectively. At six weeks, mice were anesthetized, exsanguinated, and major organs collected for processing as above.

### Serology

At the time of animal sacrifice, 1 mL of blood was removed using a needle in the vena cava prior to exsanguination and major organ harvest. Plasma was stored at -80°C and sent to IDEXX Bioanalytics for aspartate aminotransferase (AST), alanine aminotransferase (ALT), alkaline phosphatase (ALP), total bilirubin, blood urea nitrogen (BUN), creatinine, phosphorus, creatine kinase, triglyceride, total cholesterol, high density lipoprotein (HDL), glucose, albumin, and total protein quantification.

### Tissue Injury Scoring

Three blinded scorers (JD, BBP, and ANV) were instructed on how to perform injury scoring of photomicrographs of lung, liver, kidney, and heart tissue using the rubrics in the Online Data Supplement. Tissues from KF4 or IgG treated wild type mice were duplicated and de-identified and a score for each image determined. Unpublished cecal ligation and puncture images were used as positive controls. The average image scores were analyzed by treatment and reviewer agreement assessed.

### Lung Tissue Elastase Assay

Mouse lung elastase activity was quantified using N-Succinyl-Ala-Ala-Ala-pNitroanilide, substrate for Human Leucocyte Elastase and Porcine Pancreas Elastase (Elastin Products Company #NS945). Lung tissue was homogenized in PBS and 10 mg incubated in the assay for four hours per manufacturer instructions with KF4 or IgG antibody. Change in absorbance at 410 nm between zero and four hours was used as a measure of elastase activity.

### Statistical Analysis

Ordinal data was analyzed using Kruskal-Wallis test with Dunn’s post hoc comparison. Parametric mean linear intercept data was compared by one-way ANOVA with Holm-Sidak post hoc test. All tests were two-sided and a p-value of less than 0.05 was considered statistically significant. Parametric data is presented as line and whisker plots with lines representing mean and whiskers standard deviation. Non-parametric and ordinal data is presented as box plots. Interrater reliability measurements are presented as Kirppendorff’s alpha values defining excellent agreement as >0.8, good 0.67 to 0.8, moderate 0.5-0.67, and poor <0.5. The R statistical computing platform, rstatix, irr, and ggpubr packages were used for analysis and figure generation (12–15).

## Results

### Expression of Cela1 in Murine Tissues

To understand the potential toxicities of CELA1 inhibition, we first evaluated normalized mRNA counts in the mouse ENCODE transcriptome library using NCBI Gene. Duodenum, small intestine, colon, and spleen samples had the highest count values (Figure 1A). Notably, pancreas was not in the list of evaluated tissues. We then performed immunohistochemistry on selected mouse tissues. As expected, expression was strong throughout the pancreas (Figure 1B). In the kidney, a subset of cells that appeared to be distal convoluted tubules had Cela1 protein (Figure 1C). In the small intestine, Cela1 protein was present in enterocytes in the luminal 1/3 of the villus. This staining was intracellular which with that and mRNA expression indicated that this was not from pancreas-secreted Cela1 (Figure 1D). To further identify cell types potentially expressing Cela1 in the spleen and in consideration of our previous finding that there were more numerous leukocytes in the lungs of *Cela1^-/-^* mice (8), we evaluated what leukocytes were reported has having CELA1 mRNA in the Human Protein Atlas (16). The two cell types with the highest mRNA levels were basophils and Naïve CD4 T-cells (Figure 1E). Consistent with *Cela1* expression in Naïve CD4 T-cells, we observed Cela1 staining in the red pulp of the spleen between follicles (Figure 1F). In the mouse lung, Cela1 was expressed in sparse, contained cells (Figure 1G) consistent with its expression in alveolar type 2 cells. These data indicated that Cela1 inhibition could result in digestive, renal, or immunological toxicities.

**Figure 1:**
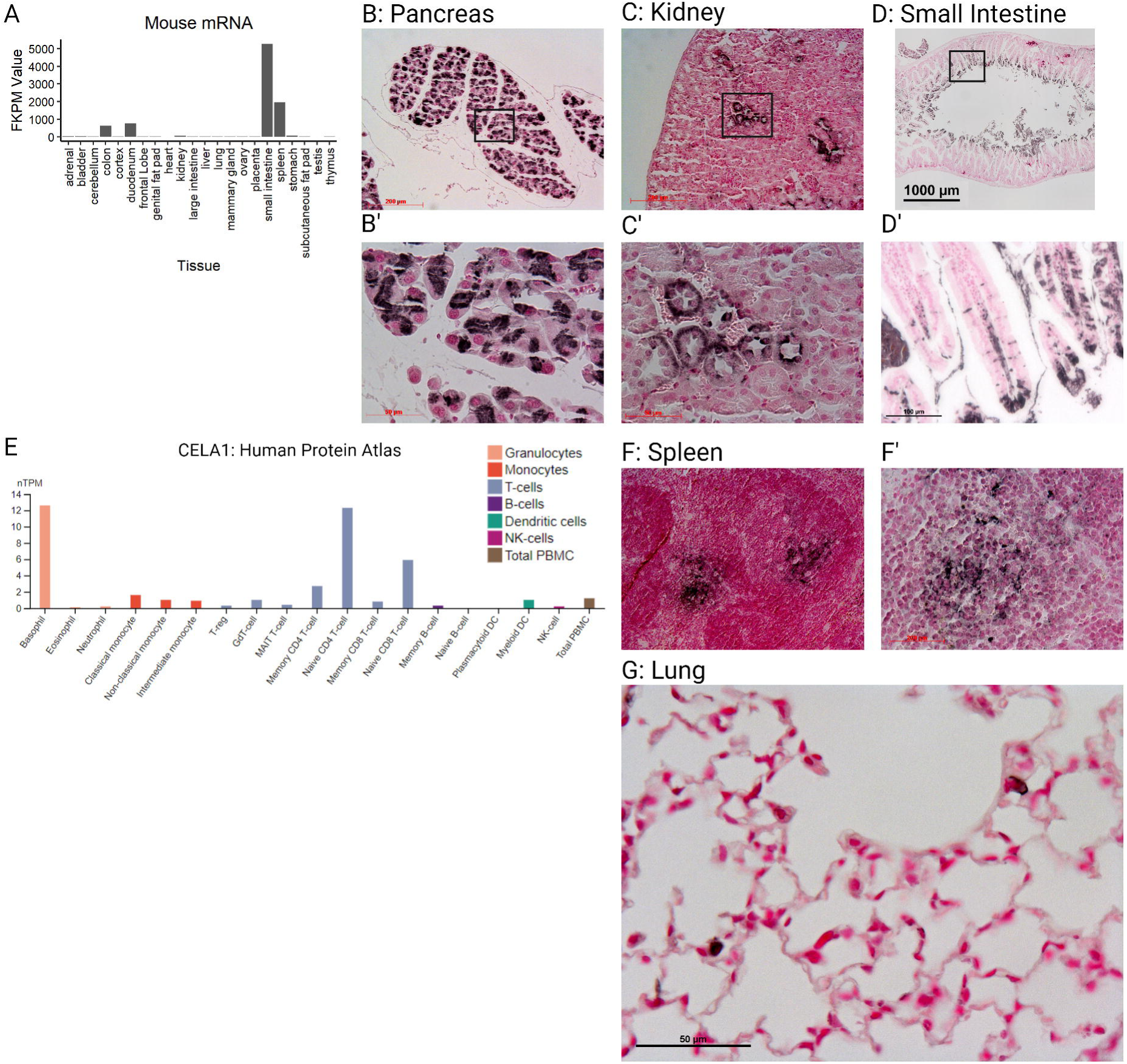
Expression of Chymotrypsin-like Elastase 1 in Mouse Tissues. (A) Assessment of mRNA levels of various mouse tissues showed that the highest *Cela1* mRNA levels were in the small intestine, spleen, and kidney. FKPM = Fragments per kilobase per Million mapped reads. (B) Pancreas tissue was not included in this database, but as a pancreatic protease, Cela1 was expressed in pancreatic acinar cells. Scale bar = 200 µm (B’) Higher magnification of B. (C) In the adult mouse kidney, Cela1 was present in a subset of tubules that were in a location and with morphology consistent with distal convoluted tubules. Scale bar 50 µm. (C’) Higher magnification of C. (D) Jejunum from the mouse showed staining in cells and/or debris in the lumen and in the luminal quarter of the villus. Scale bar = 1000 µm. (D’) Magnifying the distal villus showed intracellular staining of enterocytes. (E) To identify what types of cells might be expressing Cela1 in the spleen, we used a query tool in the Human Protein Atlas which indicated highest expression in basophils and Naïve CD4 T-cells. nTPM = normalized transcripts per million. (F) In the spleen, Cela1 staining was seen between follicles which is consistent with Naïve CD4 T-cell location. (F’) higher magnification of F. (G) In the mouse lung, rare Cela1-expressing, rounded cells are observed in alveolar walls. Scale bar = 50 µm. Created with BioRender.com.

### Toxicity Assessment of KF4 anti-CELA1 Antibody

To test for toxic effects of KF4, we assessed administered intraperitoneal injections totaling 5 mg/kg KF4, 12.5 mg/kg KF4, 25 mg/kg KF4 or 25 mg/kg KF4 weekly for four weeks. We recorded mouse health assessments three times weekly, mouse mass weekly, and at four weeks collected blood for serology and performed blinded tissue injury scoring of mice. Throughout the four weeks, no mouse showed evidence of distress (i.e. scored less than 3 on a 0-3 scale of mouse health). Mice treated with 25 mg/kg KF4 per week had less weight gain than lower dose KF4 or IgG treatment (Figure 2A). Serology of these mice after 4 weeks of treatment showed no differences in liver function tests, kidney function tests, myocardial or other muscle injury, or lipid profile except for a slightly higher blood urea nitrogen (BUN) level in KF5 25 mg/kg treated mice (Figure 2B). These data suggest mild toxicity in mice treated with KF4 25 mg/kg weekly.

**Figure 2:**
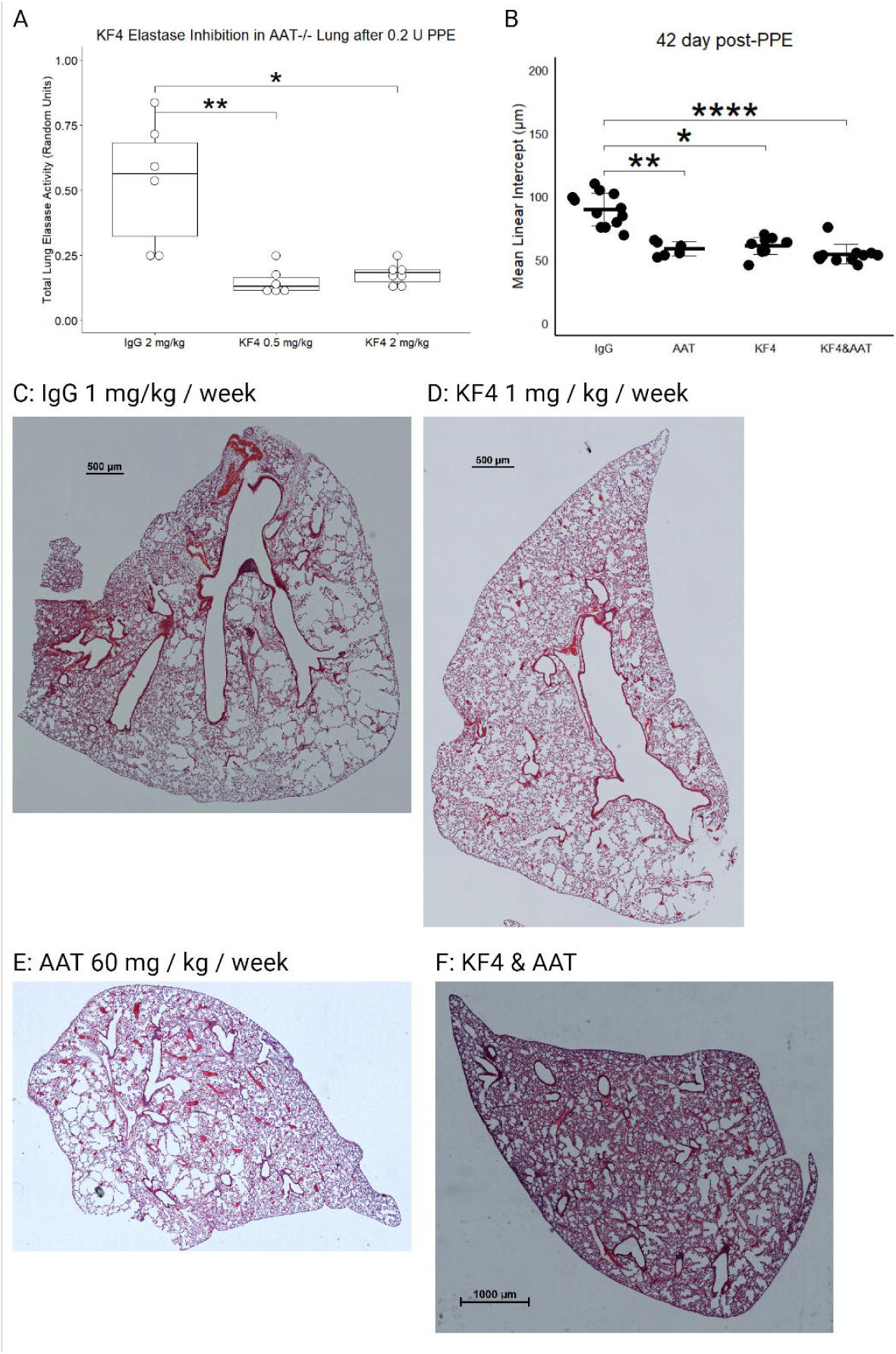
Mouse Mass and Serology Toxicity Assessments. (A) Four weeks of treatment with KF4 25 mg/kg resulted in slower weight gain compared with 5 and 12.5 mg/kg KF4 and IgG 25 mg/kg. * p<0.01, ** p<0.01, #p=0.1 KF4 25 mg/kg compared to KF4 5 mg/kg. (B) Serology from mice identified no differences in aspartate aminotransferase (AST), alanine aminotransferase (ALT), alkaline phosphatase (ALP), total bilirubin, creatinine, phosphorus, creatine kinase, triglyceride, total cholesterol, high density lipoprotein (HDL), glucose, albumin, or total protein. Blood urea nitrogen (BUN) was significantly higher in KF4 25 mg/kg compared to KF4 12.5 mg/kg treated mice (p<0.05). Created with BioRender.com.

We performed three-reviewer blinded histological scoring of lung, liver, kidney, and heart tissues from these mice. As a positive control, we used tissues from mice subjected to cecal ligation and puncture (CLP)—a mouse sepsis model in which these scoring systems have been validated. Sham treated mice in these experiments were also evaluated. For antibody-treated and control images, three blinded reviewers scored each image on four criteria using the rubrics in the Online Data Supplement.

In histologic evaluation of lung injury, both CLP and sham animals had more evidence of lung injury than antibody-treated animals (Figure 3A). Reviewer 1 consistently scored animals has having a higher level of lung injury resulting in a Krippendorff’s alpha coefficient of 0.35, but this difference was similar across groups. In otherwise healthy mice, escalating doses of KF4 do not cause any greater degree of lung injury than that of non-specific immunoglobulin and levels similar to that of an animal undergoing a sham abdominal surgery.

**Figure 3:**
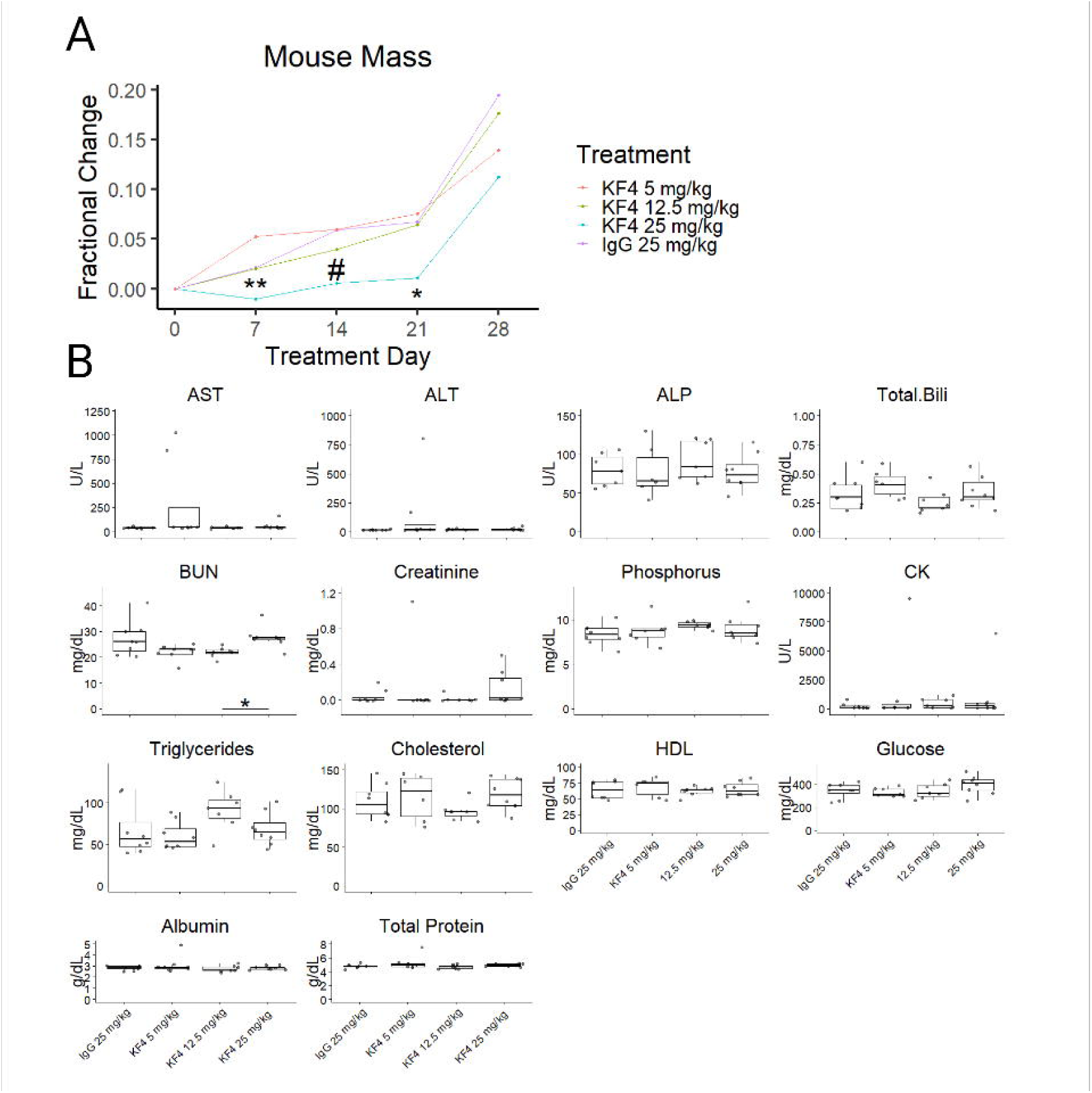
Tissue Injury Scoring. (A) Lung injury was less in KF4 and IgG treatment groups compared to lung from animals following cecal ligation and puncture (CLP). (B) Liver injury was also lower in antibody treatment groups but there was increased injury in KF4 25 mg/kg and IgG 25 mg/kg compared to KF4 5 mg/kg. (C) Kidney injury was less with KF4 or IgG treatment compared to sham and CLP animals. (D) Myocardial injury scores were elevated in CLP compared to KF4 12.5 mg/kg and IgG 25 mg/kg treatments, but the absolute difference was small. *p<0.05, **p<0.01, ***p<0.001, ****p<0.00001 by Dunn’s post hoc test. Created with BioRender.com.

A similar pattern was observed in the liver. The injury levels of both CLP-treated animals had higher injury levels than all other groups, and sham-treated animals had greater injury than KF4 5 mg/kg and 12.5 mg/kg doses. A small but significant increase in liver injury was noted comparing both IgG and KF4 25 mg/kg doses compared to KF4 5 mg/kg (Figure 3B). Inter-reviewer agreement was moderate with a Krippendorff’s alpha of 0.54.

For kidney analysis, CLP and sham operated animals had higher levels of injury than any of the antibody-treatedgroups (Figure 3C). There was substantial interrater variability, and agreement was poor with an alpha value of 0.32.

For myocardial injury scoring, only mild injury was noted in the CLP group which was greater than the injury observed in KF4 12.5 mg/kg and IgG 25 mg/kg administration (Figure 3D). Interrater agreement was again poor with an alpha of 0.279.

Taken together these data show evidence of mild liver injury with administration of KF4 or IgG 25 mg/kg that did not result in liver function test abnormalities.

### Efficacy of KF4 in Preventing Emphysema

Using the previously described low-dose PPE model of AAT-deficient emphysema (7), we performed a dose titration experiment using lung homogenate from *AAT^-/-^* mice administered 0.2 units of tracheal PPE and treated with IgG 2 mg/kg, KF4 2 mg/kg or KF4 0.5 mg/kg in a colorimetric elastase activity assay. Both KF4 doses significantly reduced lung elastase activity (Figure 4A). We then performed an efficacy assay by administering the same tracheal PPE dose to *AAT^-/-^* mice and treating them with IgG 1 mg/kg/week, KF4 1 mg/kg/week, purified human AAT 60 mg/kg/week, or combined KF4 and AAT. These last three treatment groups had less emphysema at 42 days than IgG treatment (Figure 4B). In evaluating tile scanned lung sections, it appeared that this protection was largely due to a reduction in sub-pleural emphysema (Figure 4C-F).

**Figure 4:**
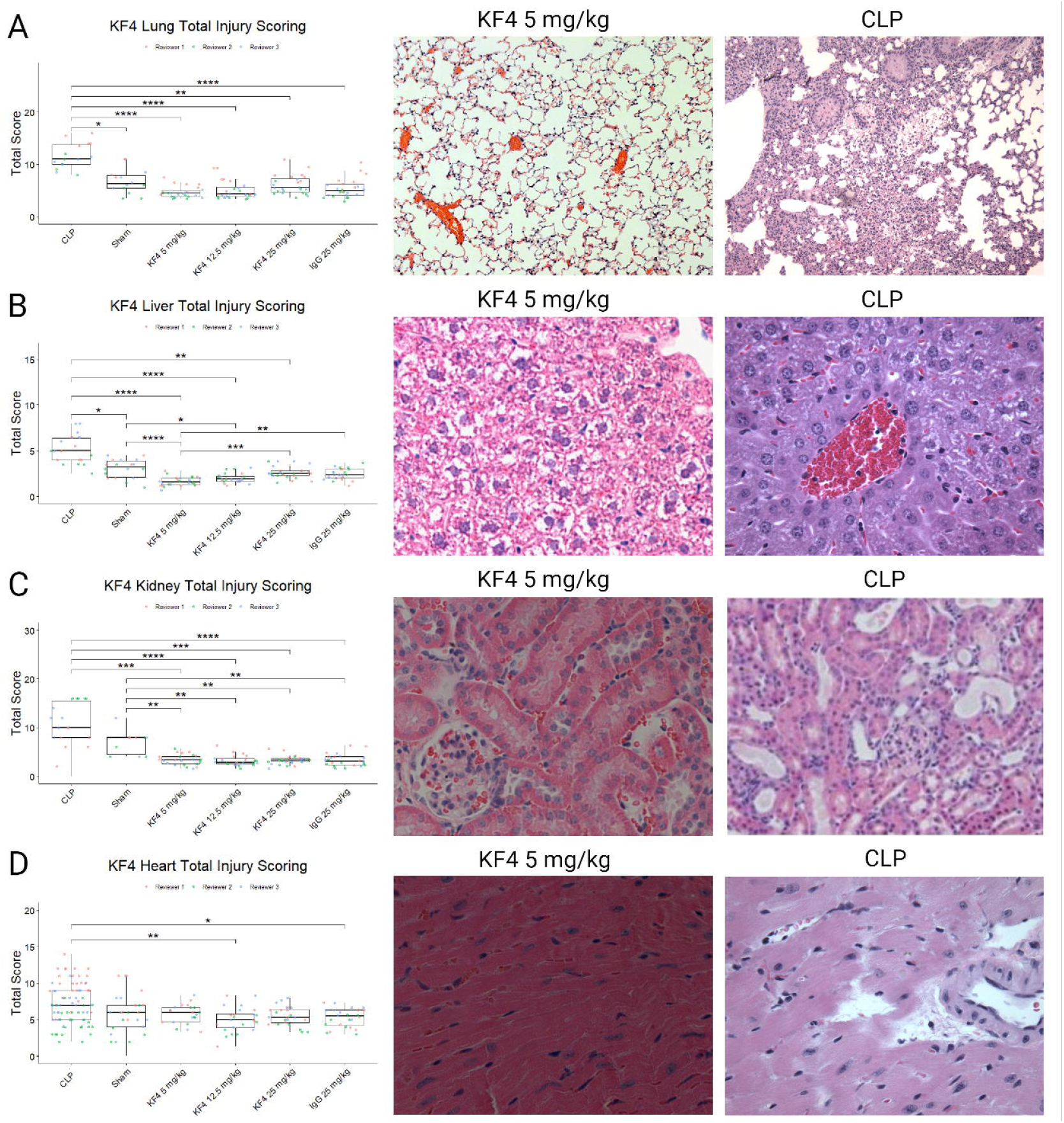
KF4 Treatment in AAT-deficient Emphysema Model. (A) At 21 days after 0.2 units of tracheal porcine pancreatic elastase, the lungs of *AAT^-/-^* mice treated with weekly KF4 0.5 mg/kg and KF4 2 mg/kg had less total elastase activity than those treated with IgG 2 mg/kg. *p<0.05, **p<0.01 by Dunn’s post hoc test. (B) Treating *AAT^-/-^*mice exposed to 0.2 units of tracheal PPE with IgG 1 mg/kg, KF4 1 mg/kg, purified human AAT 60 mg/kg, or combined AAT and KF4 showed less emphysema in the latter three groups compared to the first. *p<0.05, **p<0.01, **** p<0.0001 by Dunn’s post hoc test. (C) Tile scanned middle lobe lung section of an IgG-treated mouse showing substantial emphysema. Scale bar = 500 µm. (D) KF4-treatement had less emphysema as did (E) AAT treatment. (F) Combined treatment also had less emphysema, and in each of these there was a lack of sub-pleural airspace simplification. Created with BioRender.com.

## Discussion

In this study, we showed that the KF4 anti-CELA1 antibody and purified human AAT were similarly effective at preventing emphysema in a mouse model of AAT-deficient emphysema and that KF4 has the potential for renal and hepatic toxicity. Combining AAT and KF4 treatment did not result in less emphysema than either treatment alone. These data are supportive of a central role of CELA1 in AAT-deficient emphysema and suggest that selective CELA1 neutralization could be an alternative for purified AAT replacement therapy in AAT-deficient emphysema.

Our murine toxicity studies suggest that KF4 has a therapeutic index of at least 12.5. Renal toxicity was evidenced by an elevation of blood urea nitrogen, and evidence of histologic liver injury was observed in the highest administered KF4 dose. Due to the asynchronous nature of our experiment, we were unable to perform any hematologic assessments which is a limitation of our study. It is unclear whether reduced mouse weight with KF4 25 mg/kg treatment is due to this toxicity or intestinal malabsorption due to inhibition of pancreatic and intestinal Cela1. Interestingly, in humans *CELA1* is not expressed in pancreatic acinar cells due to a mutation in *Pancreatic Transcription Factor 1* (17, 18). This coupled with the fact that CELA1 but not CELA2A, CELA2B, CELA3A, or CELA3B is highly conserved in placental mammals (6) and loss of function mutations are more rare in CELA1 than expected (8) suggest an important, non-digestive role for CELA1. In our short-term murine toxicity studies and inhibition studies in emphysema models, at least one of these roles appears to be postnatal lung remodeling with potential intestinal and immune roles that need to be better defined.

Our findings are most consistent with our previously described model that CELA1 acts in the progression but not the initiation of emphysema (8). This would explain seemingly contradictory findings that *Cela1^-/-^* and KF4-treated mice develop less emphysema in response to 6 months of cigarette smoke exposure compared to wild-type or IgG-treated mice, (8) but cigarette smoke-exposed *AAT^-/-^&Cela1^-/-^* mice have slightly but significantly greater emphysema than *AAT^-/-^* mice (7). Daily cigarette smoke exposure causes persistent inflammation and the activity of leukocyte-associated proteases like neutrophil elastase, cathepsin G, and matrix metalloproteinases are likely greater in this model than in the PPE model used in this study. Since *AAT^-/-^* mice lack the canonical anti-protease to counter these proteases, they are perhaps more important than Cela1-mediated remodeling in the context of continued inflammation. In wild type mice, endogenous AAT mitigates the activity of these proteases, and the Cela1-effect is more prominent. Exactly how the genetic absence of *Cela1* potentiates the effect of these other proteases, whether and how Cela1 changes the behavior of Naïve CD4 T-cells, and whether these findings also occur with Cela1 inhibition need further study.

We previously reported that *Cela1^-/-^* mice gain weight at rates comparable to wild type mice (6). The finding of slower weight gain in mice treated with KF4 25 mg/kg compared to IgG 25 mg/kg suggests the possibility that other digestive enzymes are upregulated with Cela1 is absent throughout development; although, we cannot rule out some other developmental effect. The feces of mice treated with KF4 25 mg/kg were indistinguishable from those in other groups, but a biochemical assessment of those feces might reveal important differences.

There are several important limitations of this study. First, we did not evaluate long-term toxicity. The fact that all mice gained weight in this study is reassuring, but it may be that longer term toxicity studies would show worsening liver injury. Of course, the dose at which injury was observed was 10-fold higher than the therapeutic dose, and it may be that KF4 is effective at even lower doses. The model of AAT-deficient emphysema used was previously shown to be progressive (7), but it is a post-injury model and lacks the clinical relevance that other models like cigarette smoke exposure have. Lastly, the combination of KF4 and AAT replacement could have caused some amount of injury associated with a higher osmotic load obscuring potential benefit of combination therapy.

In conclusion, we found that the KF4 anti-CELA1 antibody and purified human AAT were similarly effective at preventing emphysema in a mouse model of AAT-deficient emphysema but that there was no benefit to combined therapy.

## Abbreviations

AAT: Alpha-1 Antitrypsin
CELA1: Chymotrypsin-like Elastase 1
COPD: Chronic Obstructive Pulmonary Disease
MLI: Mean Linear Intercept
PPE: Porcine Pancreatic Elastase

## Acknowledgements

We would like to thank Mark Branson at the University of Florida for providing purified, recombinant human alpha-1 antitrypsin.

## Author Participation or Contribution Statement

A-Substantial contributions to the conception or design of the work

B-Substantial contribution to the acquisition, analysis, or interpretation of data

C-Drafting the work or revising it critically for important intellectual content

D-Final approval of the version to be published

E-Agreement to be accountable for all aspects of the work in ensuring that questions related to the accuracy or integrity of any part of the work are appropriately investigated and resolved.

## Data Sharing Statement

The data used in manuscript presentation is publicly available at 10.6084/m9.figshare.25403464

## References

1. Busch R, et al. Genetic Association and Risk Scores in a COPD Meta-Analysis of 16,707 Subjects. American journal of respiratory cell and molecular biology. [published online ahead of print: February 2017]. 10.1165/rcmb.2016-0331OC.

2. Gotzsche PC, Johansen HK. Intravenous alpha-1 antitrypsin augmentation therapy for treating patients with alpha-1 antitrypsin deficiency and lung disease. The Cochrane database of systematic reviews. 2016;9:Cd007851.

3. Adam Wanner MD. Towards New Therapeutic Solutions for Alpha-1 Antitrypsin Deficiency: Role of the Alpha-1 Foundation. Chronic Obstructive Pulmonary Diseases:Journal of the COPD Foundation;7(3):147–150.

4. Heinz A, et al. The action of neutrophil serine proteases on elastin and its precursor. Biochimie. 2012;94(1):192–202.

5. Guyot N, et al. Unopposed cathepsin G, neutrophil elastase, and proteinase 3 cause severe lung damage and emphysema. The American journal of pathology. 2014;184(8):2197–210.

6. Joshi R, et al. Role for Cela1 in Postnatal Lung Remodeling and AAT-deficient Emphysema. American journal of respiratory cell and molecular biology. 2018;59(2):167–178.

7. Devine AJ, et al. Chymotrypsin-like Elastase-1 Mediates Progressive Emphysema in Alpha-1 Antitrypsin Deficiency. Chronic Obstr Pulm Dis. [published online ahead of print: August 1, 2023]. 10.15326/jcopdf.2023.0416.

8. Ojha M, et al. Anti-CELA1 antibody KF4 prevents emphysema by inhibiting stretch-mediated remodeling. JCI Insight. 2024;9(1). 10.1172/jci.insight.169189.

9. Inata Y, et al. Age-dependent cardiac function during experimental sepsis: effect of pharmacological activation of AMP-activated protein kinase by AICAR. American Journal of Physiology-Heart and Circulatory Physiology. 2018;315(4):H826–H837.

10. Inata Y, et al. Autophagy and mitochondrial biogenesis impairment contribute to age-dependent liver injury in experimental sepsis: dysregulation of AMP-activated protein kinase pathway. The FASEB Journal. 2018;32(2):728–741.

11. Dunnill MS. Quantative Methods in the Study of Pulmonary Pathology. Thorax. 1962;17(4):320–328.

12. R Core Team. R: A Language and Environment for Statistical Computing. Vienna, Austria: R Foundation for Statistical Computing; 2019.

13. Kassambara A. rstatix: Pipe-Friendly Framework for Basic Statistical Tests. 2020. https://CRAN.R-project.org/package=rstatix. Accessed August 20, 2020.

14. Kassambara A. ggpubr: Publication Ready Plots - Articles - STHDA [Internet]. 2020. http://www.sthda.com/english/articles/24-ggpubr-publication-ready-plots/. Accessed May 15, 2020.

15. Gamer M, Lemon J, Fellows I. irr: Various Coefficients of Interrater Reliability and Agreement. 2019. https://CRAN.R-project.org/package=irr.

16. The Human Protein Atlas [Internet]. https://www.proteinatlas.org/. Accessed January 9, 2020.

17. Rose SD, MacDonald RJ. Evolutionary silencing of the human elastase I gene (ELA1). Human molecular genetics. 1997;6(6):897–903.

18. Rose SD, et al. A single element of the elastase I enhancer is sufficient to direct transcription selectively to the pancreas and gut. Molecular and cellular biology. 1994;14(3):2048–57.

