## Supplementary material for "KF4 anti-CELA1 Antibody and Purified α1-Antitrypsin Have Similar but Not Additive Efficacy in Preventing Emphysema in Murine α1-Antitrypsin Deficiency": Online Data Supplement-1

### Histological Signs of Liver Injury

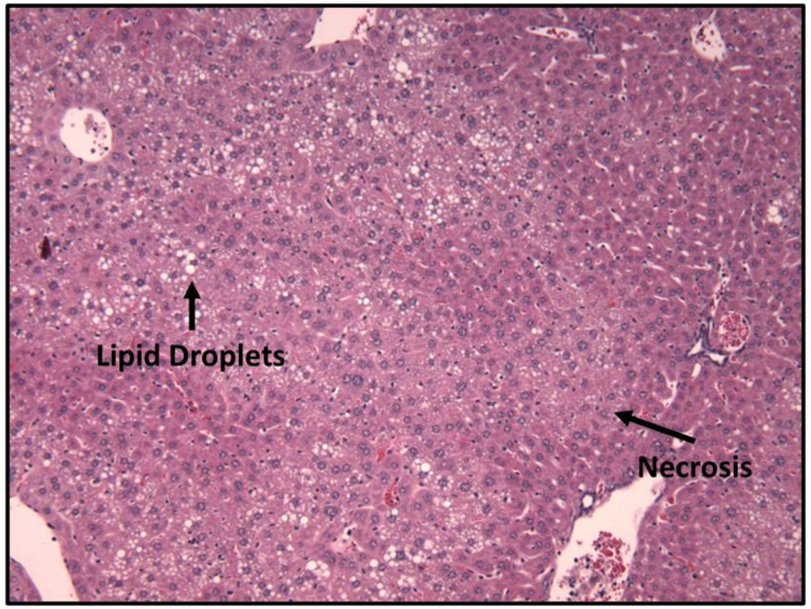


**Healthy Liver Tissue**

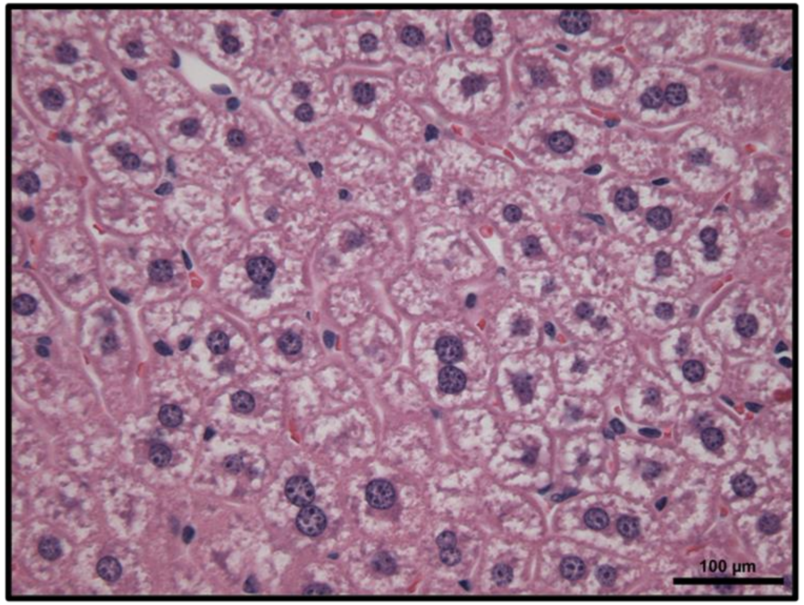


**Septic Liver Tissue**

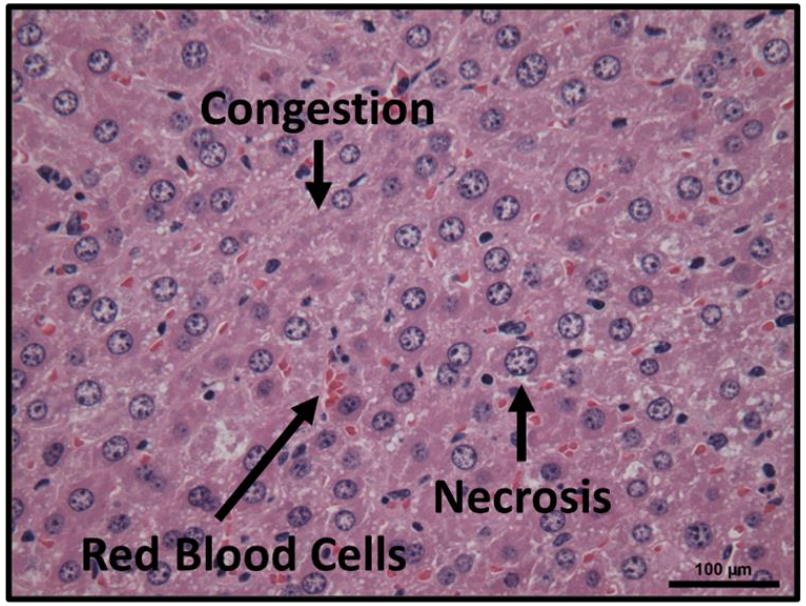

**Scores**
**0 = no injury**
**1 = minimal (0-25%of the section)**
**2 = mild (25-50%)**
**3 = significant (50-75%)**
**4 = severe (more than 75%)**

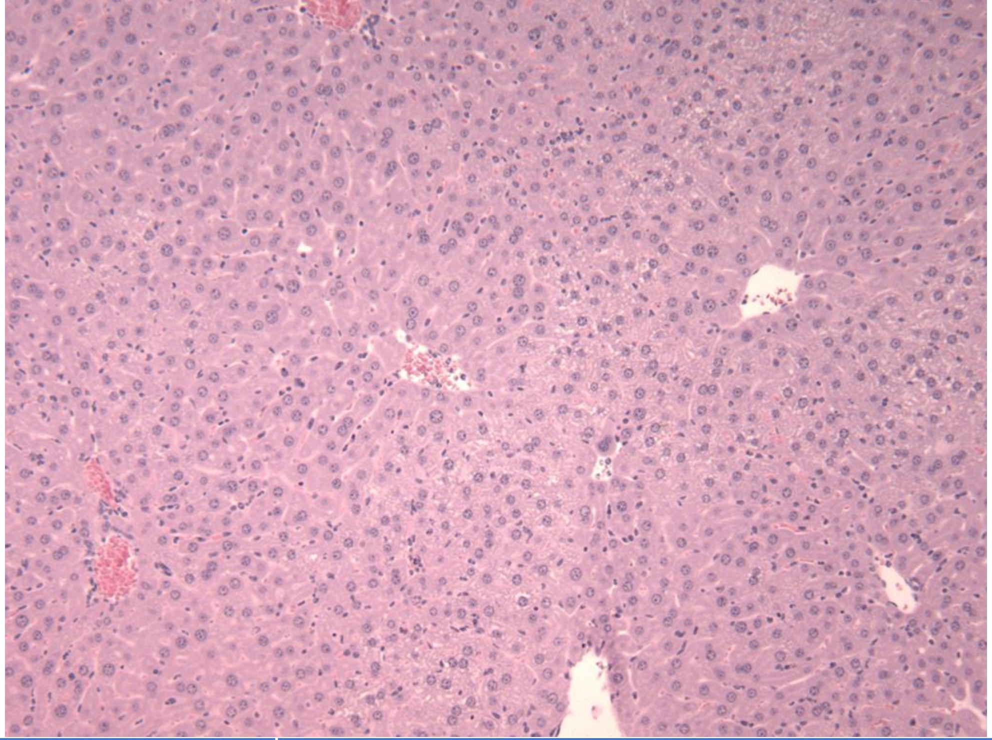

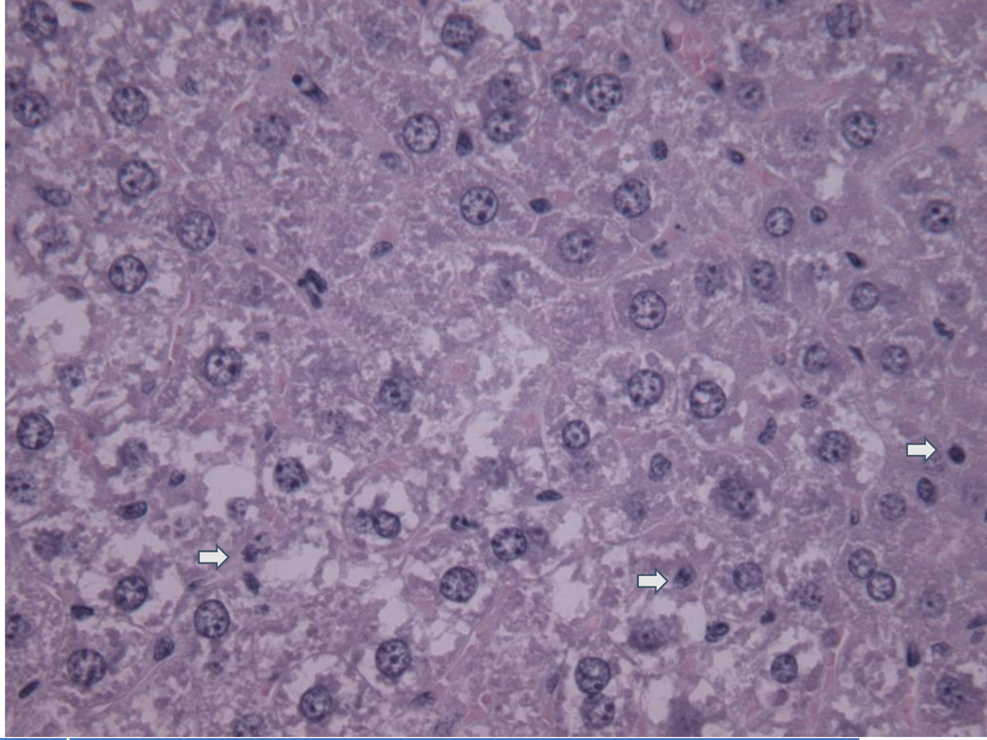

| | 10x | 40x |
|---|---|---|
| Sinusoid congestion and/or edema (10X) | 3 | |
| Infiltration of red blood and inflammatory cells (10X) | 2 | |
| Presence of lipid droplets or vacuoles (10 & 40X) | 0 | 0 |
| Necrosis (10 & 40X) | 3 | 3 |
| Total | | |

**Scores**
0 = no injury
1 = minimal (0-25%of the section)
2 = mild (25-50%)
3 = significant (50-75%)
4 = severe (more than 75%)

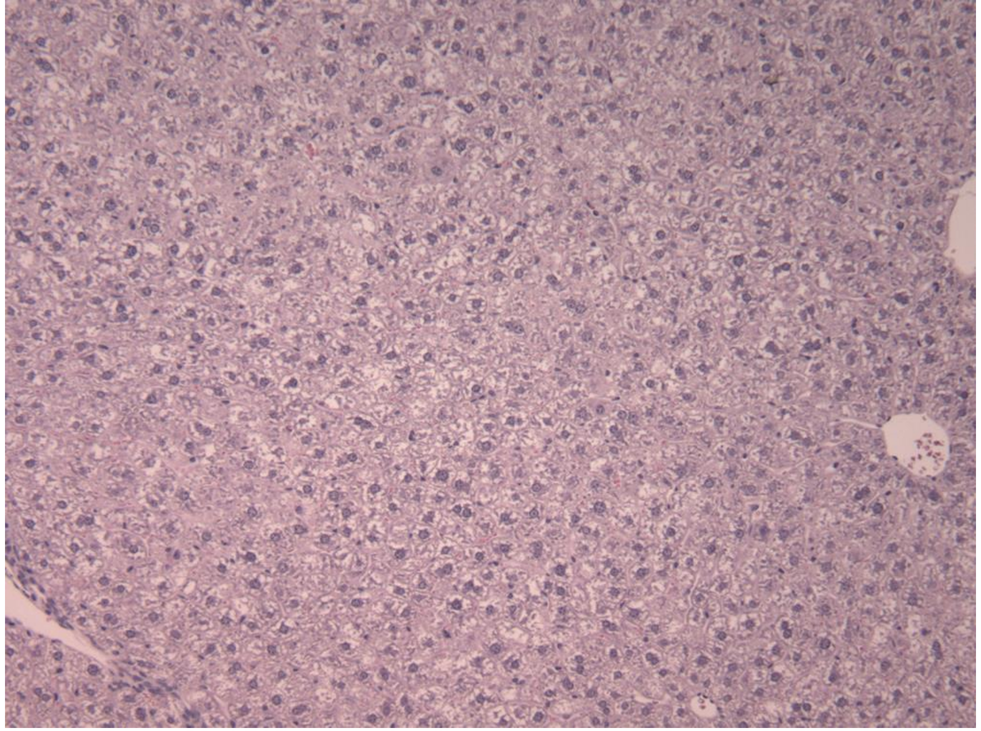

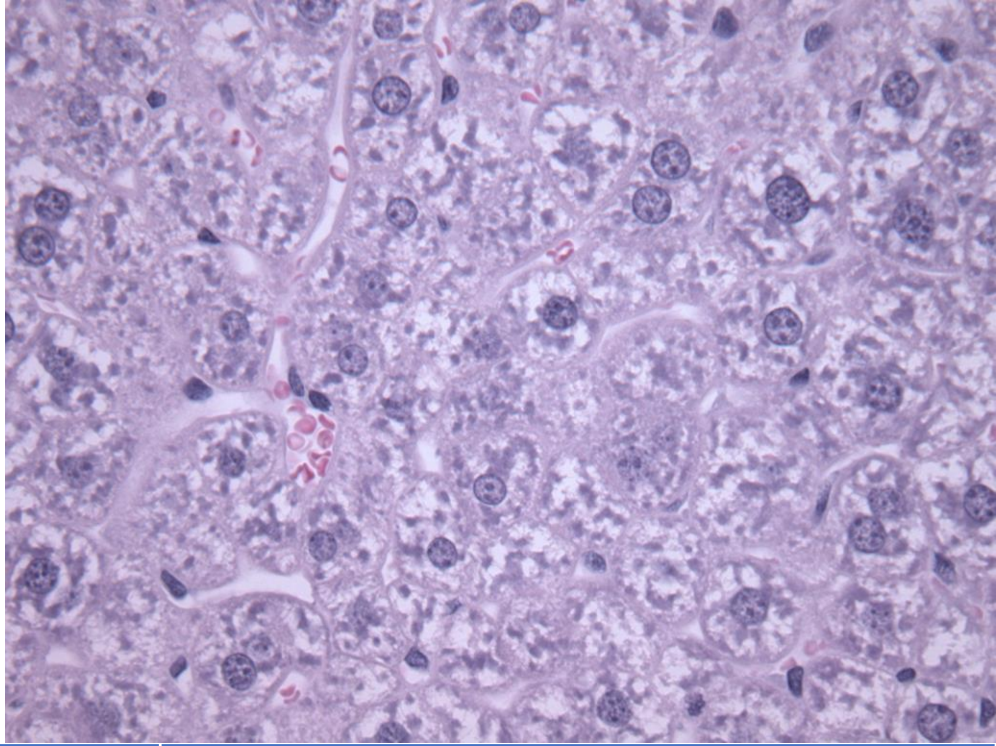

| | 10x | 40x |
|---|---|---|
| Sinusoid congestion and/or edema (10X) | 0 | |
| Infiltration of red blood and inflammatory cells (10X) | 0 | |
| Presence of lipid droplets or vacuoles (10 & 40X) | 0 | 0 |
| Necrosis (10 & 40X) | 1 | 1 |
| Total | | |

**Scores**

**0 = no injury**
**1 = minimal (0-25%of the section)**
**2 = mild (25-50%)**
**3 = significant (50-75%)**
**4 = severe (more than 75%)**

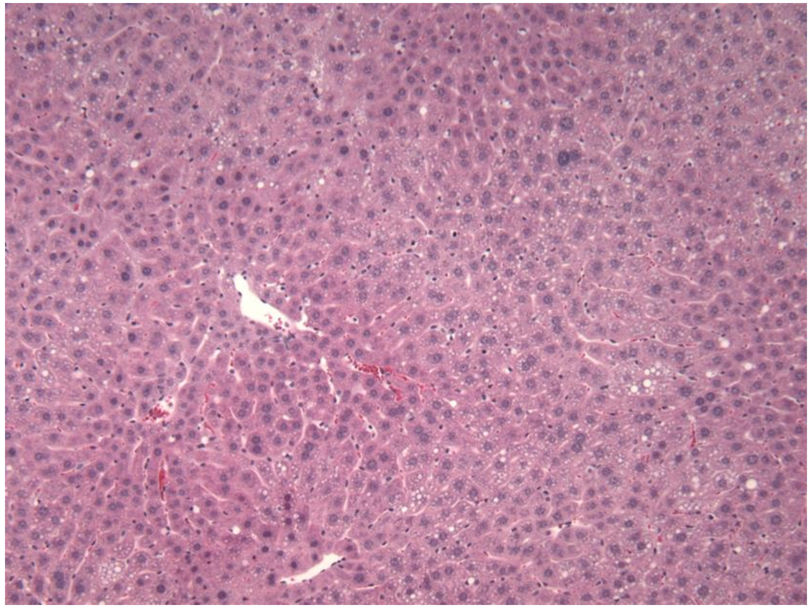

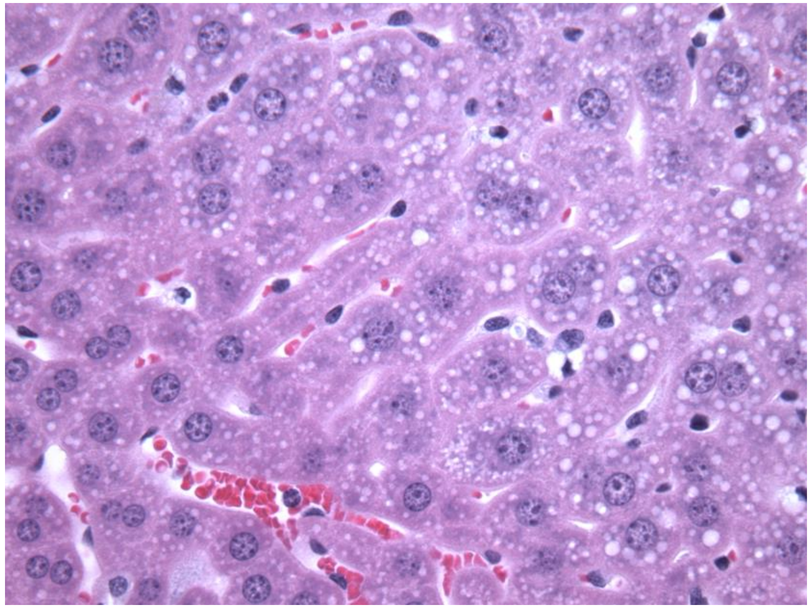

| | 10x | 40x |
|---|---|---|
| Sinusoid congestion and/or edema (10X) | 3 | |
| Infiltration of red blood and inflammatory cells (10X) | 3 | |
| Presence of lipid droplets or vacuoles (10 & 40X) | 3 | 4 |
| Necrosis (10 & 40X) | 3 | 3 |
| Total | | |
