## Supplementary material for "KF4 anti-CELA1 Antibody and Purified α1-Antitrypsin Have Similar but Not Additive Efficacy in Preventing Emphysema in Murine α1-Antitrypsin Deficiency": Online Data Supplement-2

### Histological signs of kidney injury-10X

**Healthy Kidney Tissue**

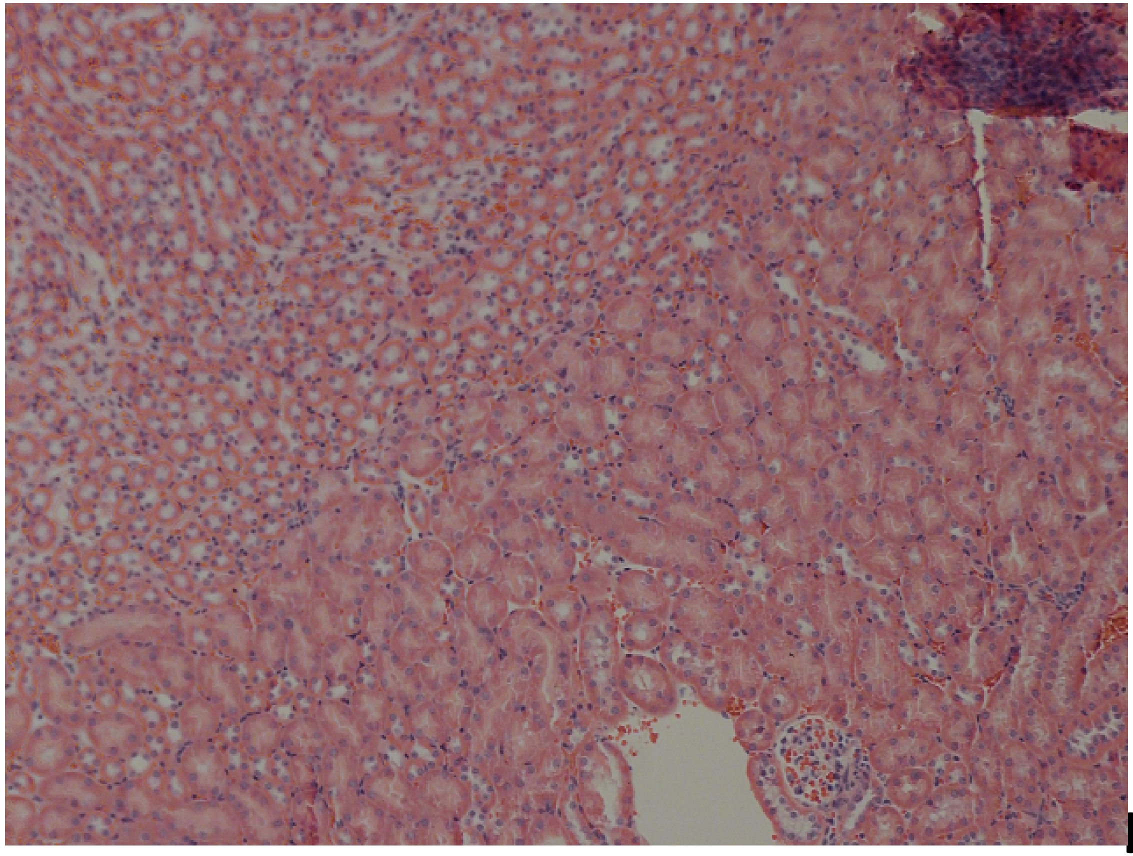

**I/R Kidney Tissue**

**Leukocytes**

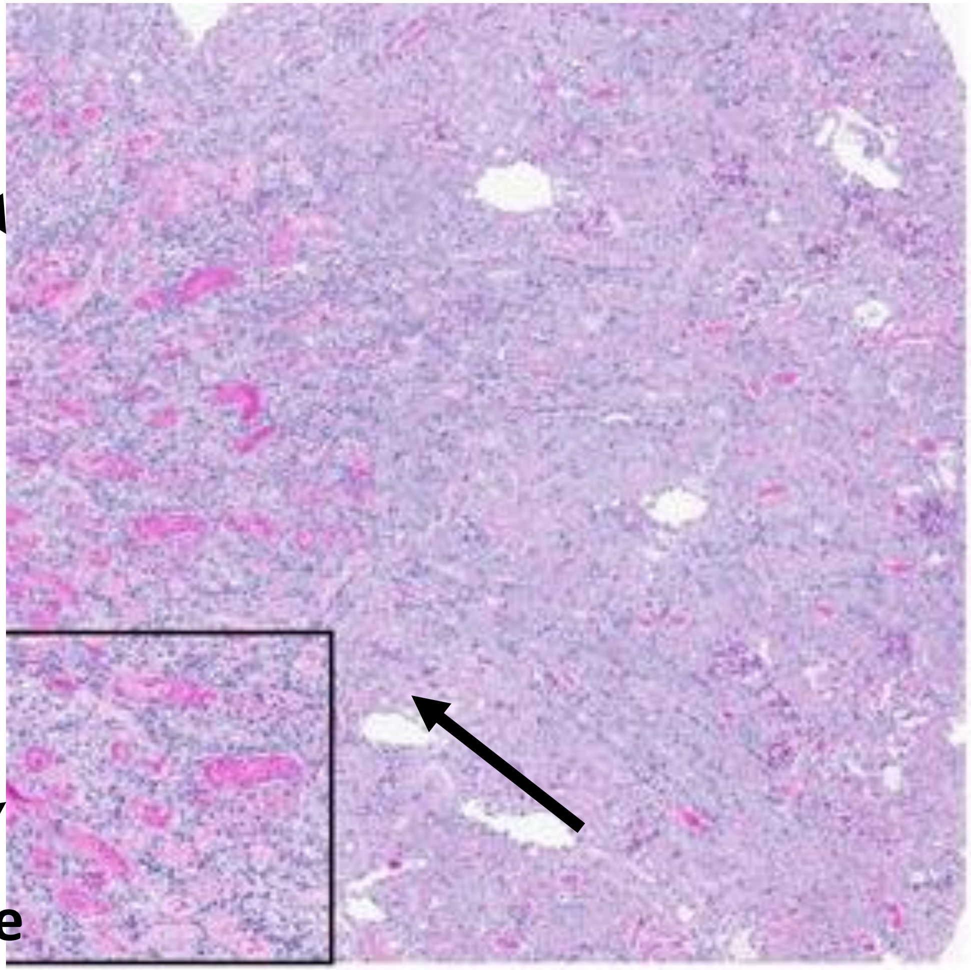

**Hemorrhage**

### Histological signs of kidney injury-40X

**Healthy Kidney Tissue**

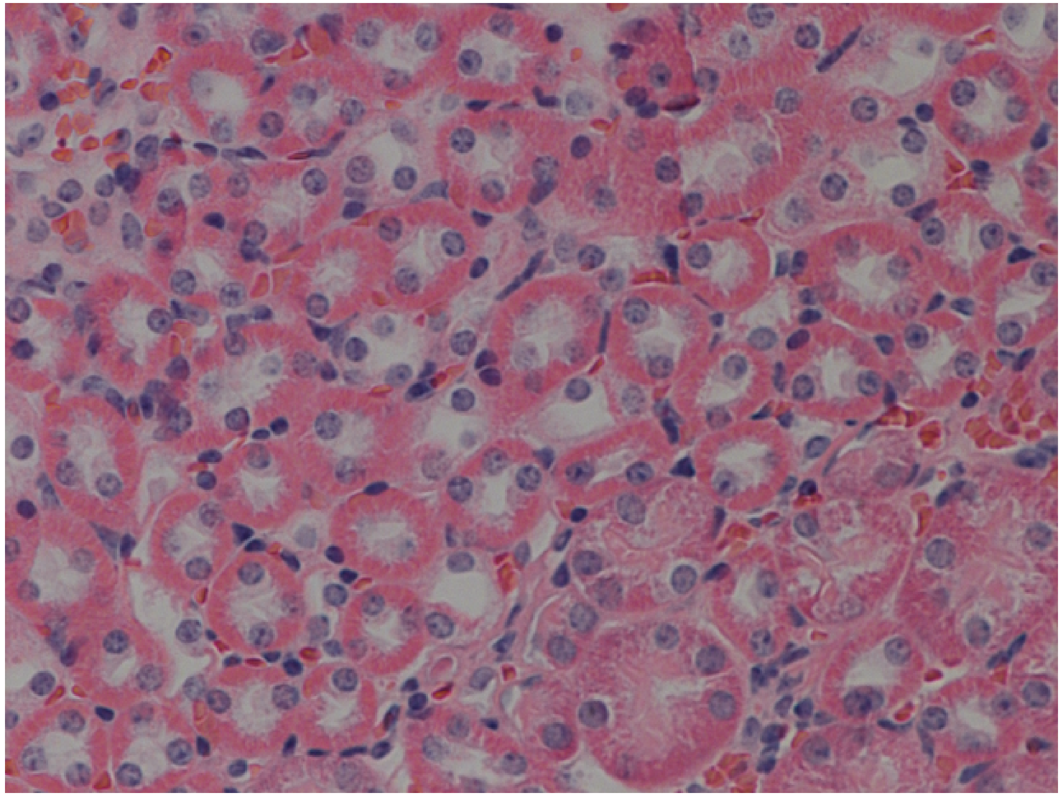

**I/R Kidney Tissue**

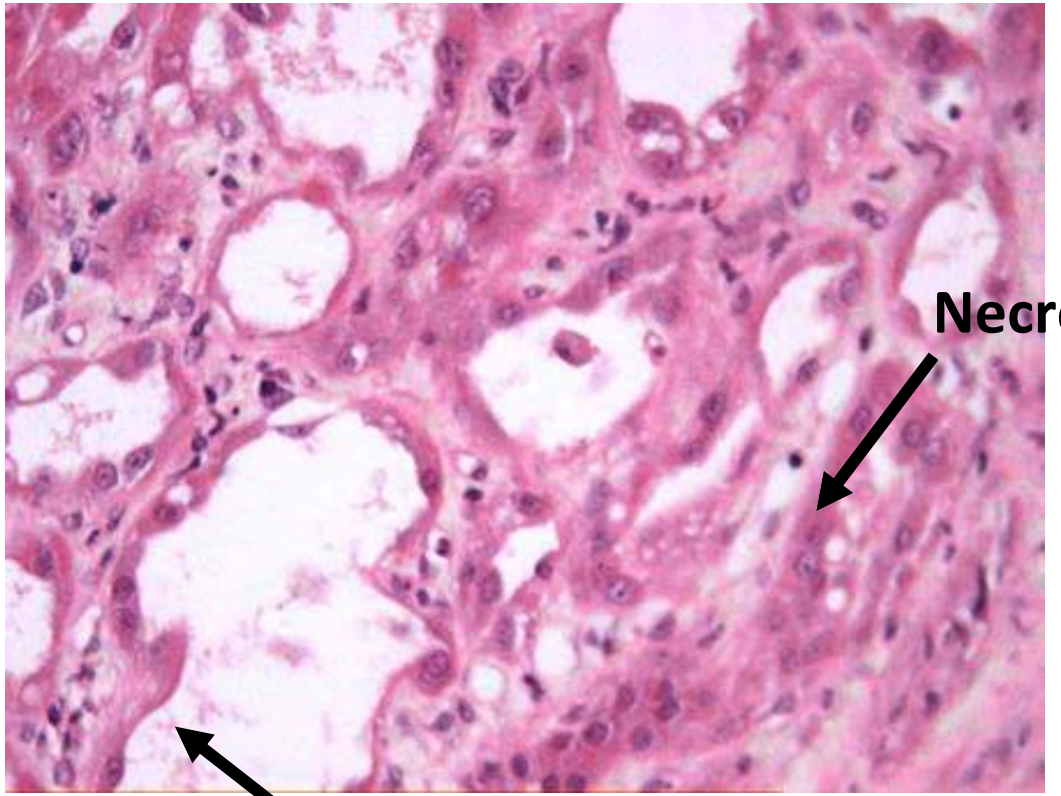

**Necrosis**

**Loss of Brush Border**

**Scores**
**0 = no injury**
**1 = minimal (0-25%of the section)**
**2 = mild (25-50%)**
**3 = significant (50-75%)**
**4 = severe (more than 75%)**

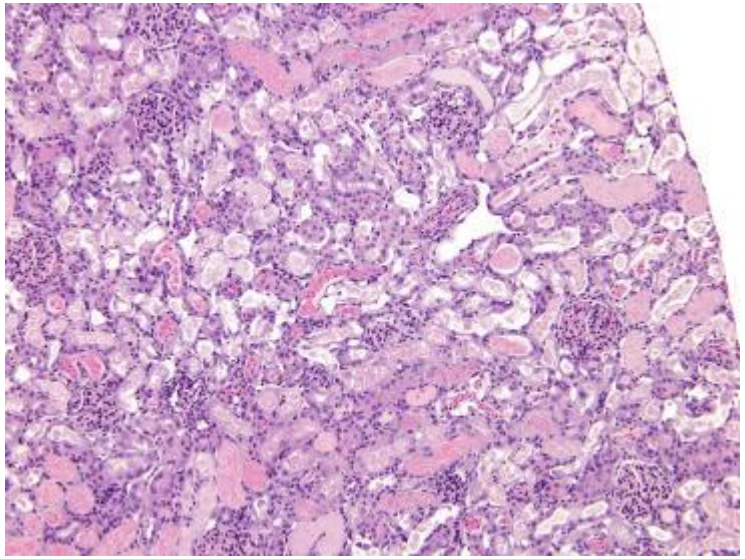

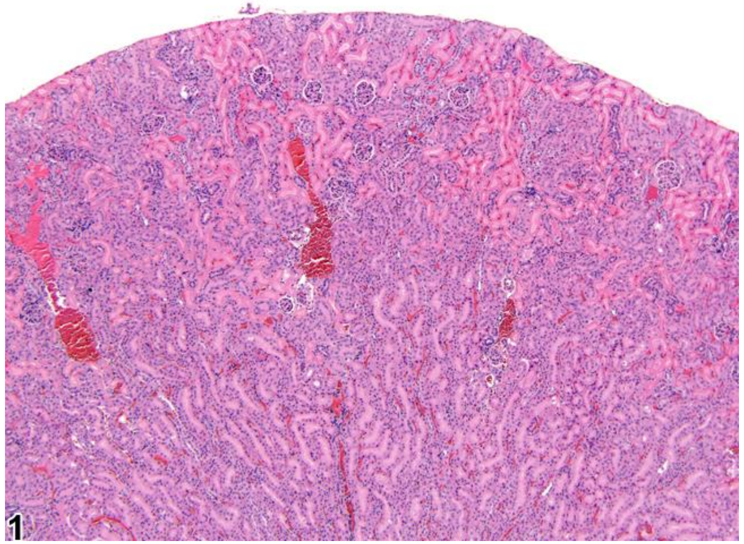


| | | |
|---|---|---|
| Loss of Brush Border (40X) | | |
| Tubular Necrosis (40X) | | |
| Neutrophil Infiltration (10X) | 3 | 4 |
| Hemorrhage/Congestion (10x) | 1 | 3 |
| Total | 4 | 7 |

**<u>Scores</u>**
**0 = no injury**
**1 = minimal (0-25%of the section)**
**2 = mild (25-50%)**
**3 = significant (50-75%)**
**4 = severe (more than 75%)**

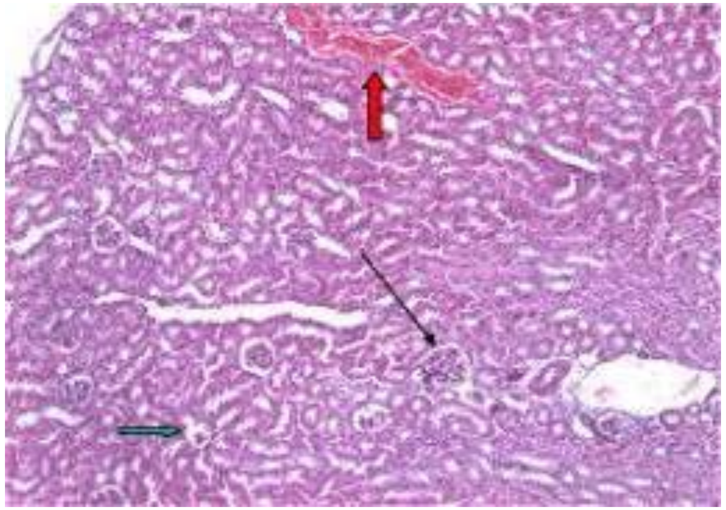

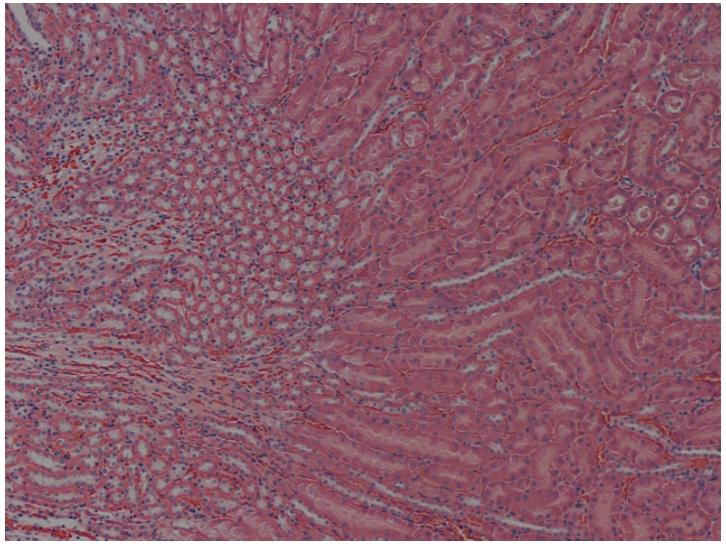

| | | |
|---|---|---|
| Loss of Brush Border (40X) | | |
| Tubular Necrosis (40X) | | |
| Neutrophil Infiltration (10X) | 0 | 0 |
| Hemorrhage/Congestion (10x) | 1 | 0 |
| Total | 1 | 0 |

### Adrenal Gland

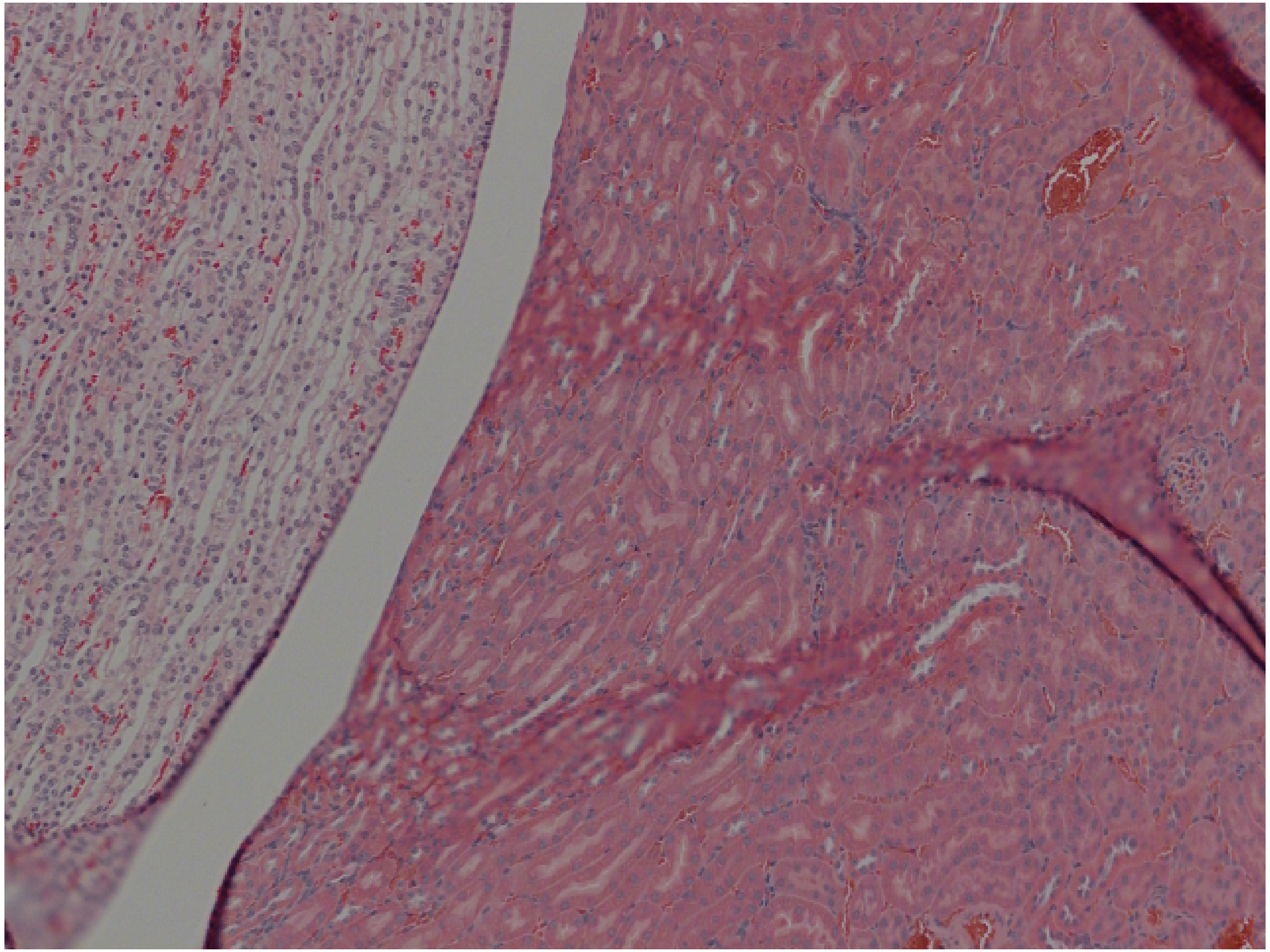

**<u>Scores</u>**
**0 = no injury**
**1 = minimal (0-25%of the section)**
**2 = mild (25-50%)**
**3 = significant (50-75%)**
**4 = severe (more than 75%)**

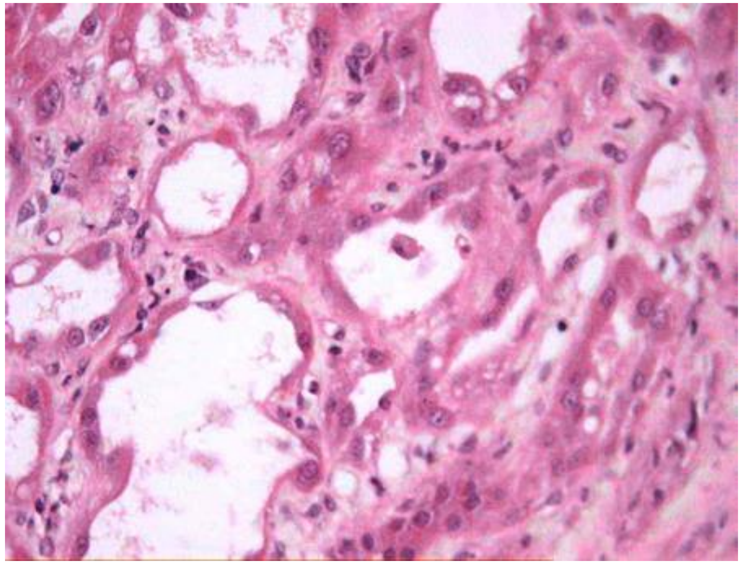

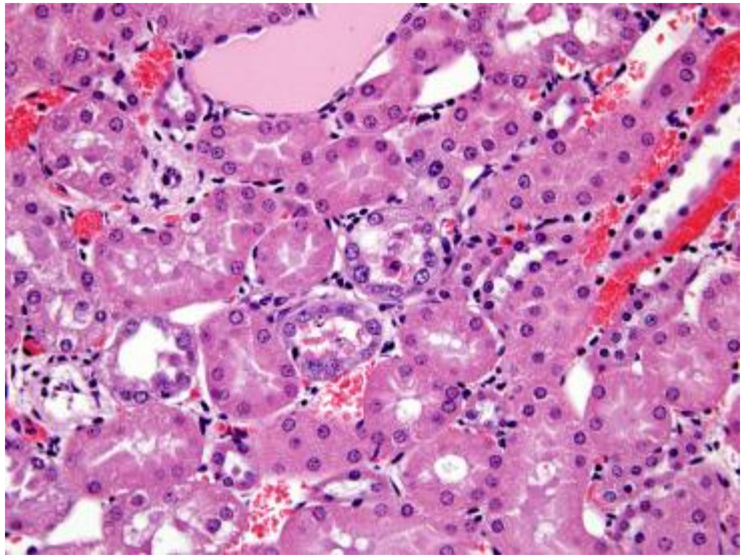

| | | |
|---|---|---|
| Loss of Brush Border (40X) | 4 | 2 |
| Tubular Necrosis (40X) | 4 | 3 |
| Neutrophil Infiltration (10X) | | |
| Hemorrhage/Congestion (10x) | | |
| Total | 8 | 5 |

**Scores**
**0 = no injury**
**1 = minimal (0-25%of the section)**
**2 = mild (25-50%)**
**3 = significant (50-75%)**
**4 = severe (more than 75%)**

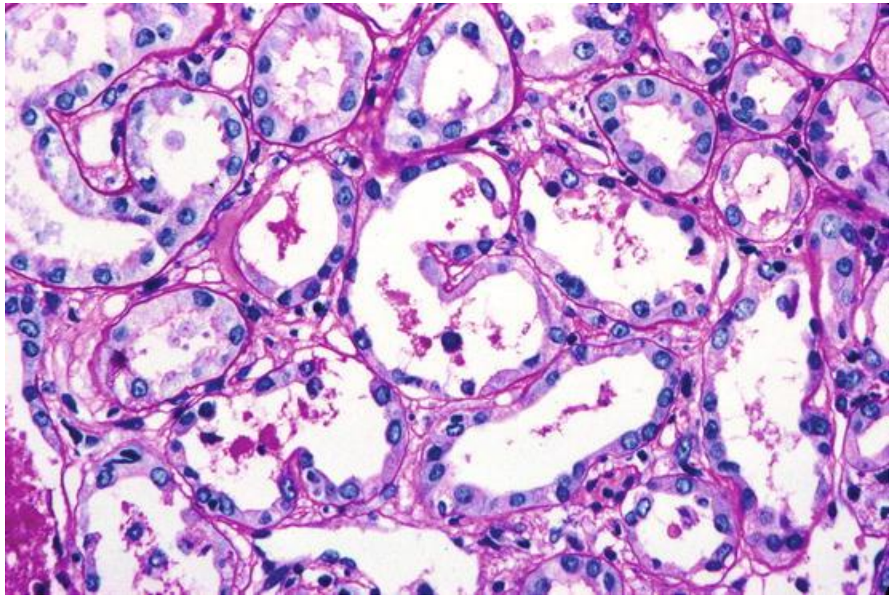

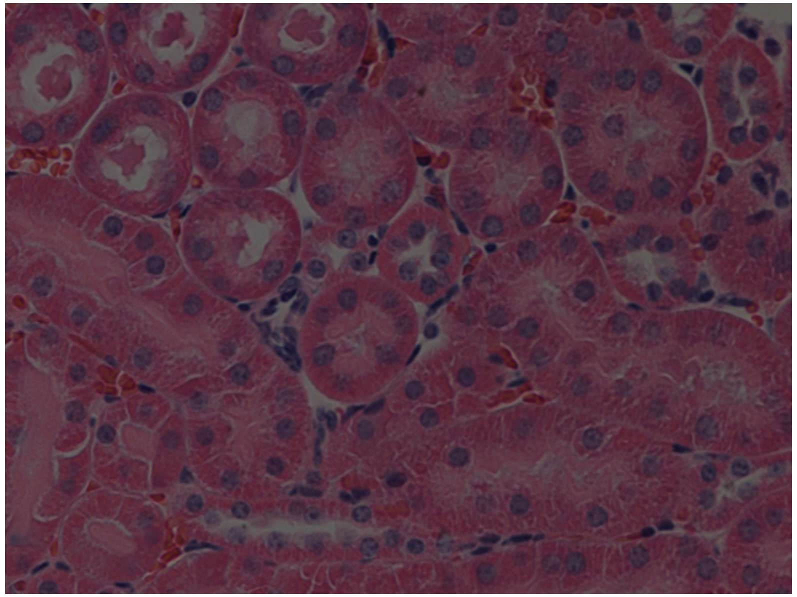

| | | |
|---|---|---|
| Loss of Brush Border (40X) | 4 | 0 |
| Tubular Necrosis (40X) | 2 | 0 |
| Neutrophil Infiltration (10X) | | |
| Hemorrhage/Congestion (10x) | | |
| Total | 8 | 0 |
