## Supplementary material for "KF4 anti-CELA1 Antibody and Purified α1-Antitrypsin Have Similar but Not Additive Efficacy in Preventing Emphysema in Murine α1-Antitrypsin Deficiency": Online Data Supplement-3

### Histological Signs of Heart Injury

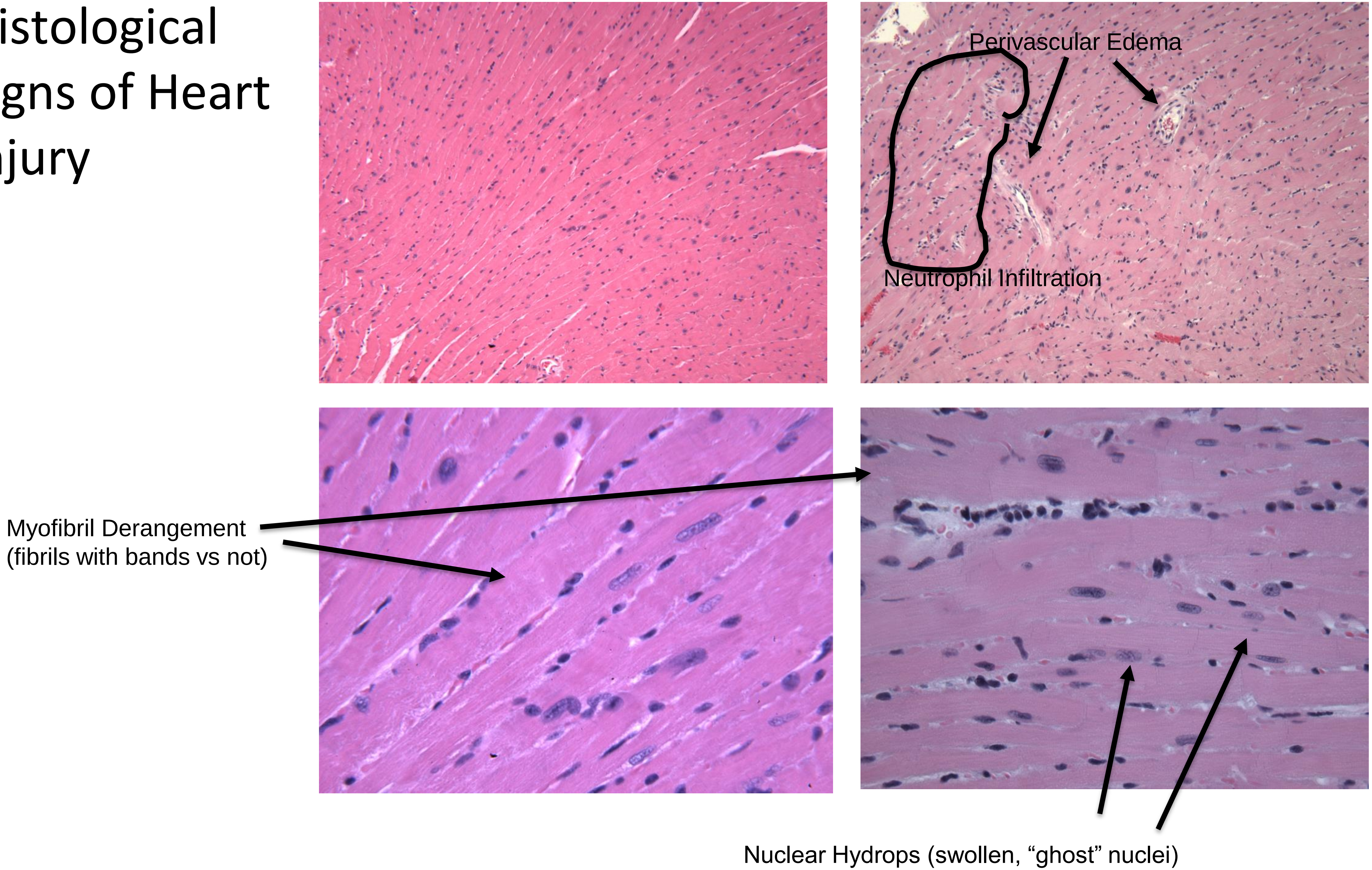

**Scores**
**0 = no injury**
**1 = minimal (0-25%of the section)**
**2 = mild (25-50%)**
**3 = significant (50-75%)**
**4 = severe (more than 75%)**

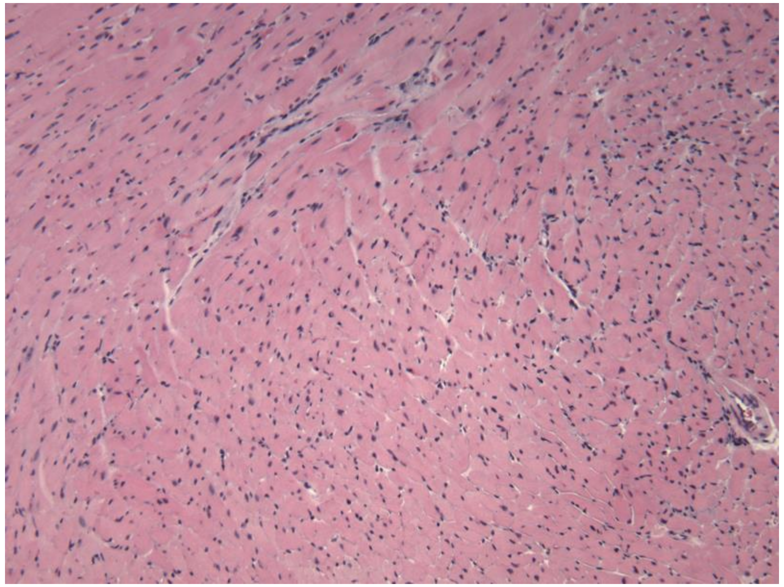

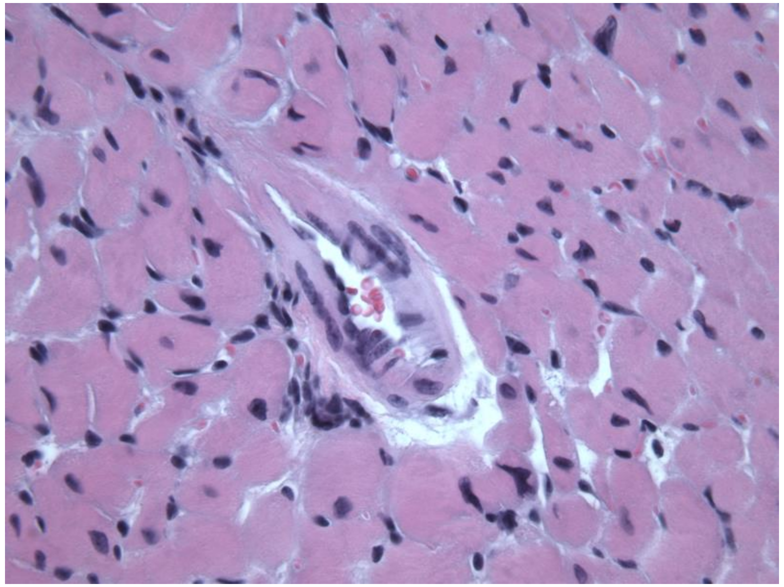

| | 10x | 40x |
|---|---|---|
| Perivascular Edema (10X) | 2 | |
| Myofibril Derangement (40X) | | 3 |
| Infiltration of Neutrophils (10X) | 3 | |
| Nuclear Hydrops (40X) | | 1 |
| Total | | |

**Scores**
**0 = no injury**
**1 = minimal (0-25%of the section)**
**2 = mild (25-50%)**
**3 = significant (50-75%)**
**4 = severe (more than 75%)**

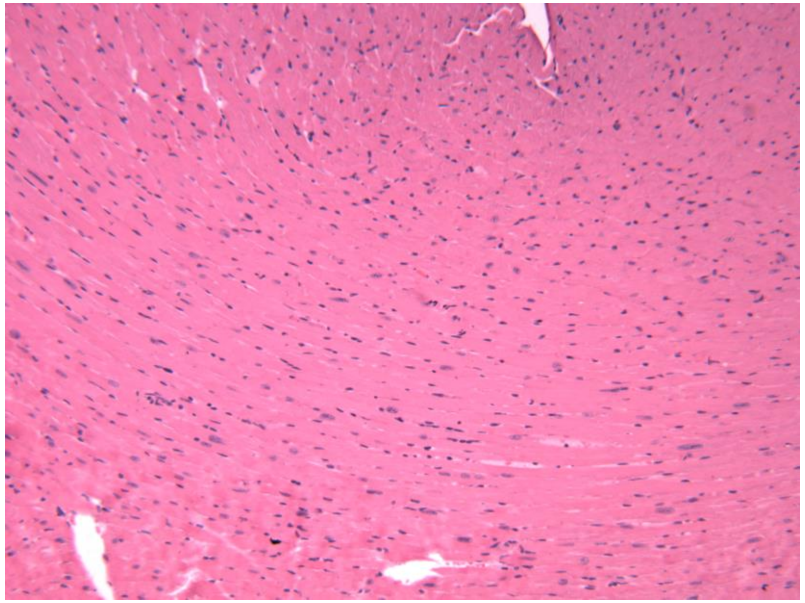

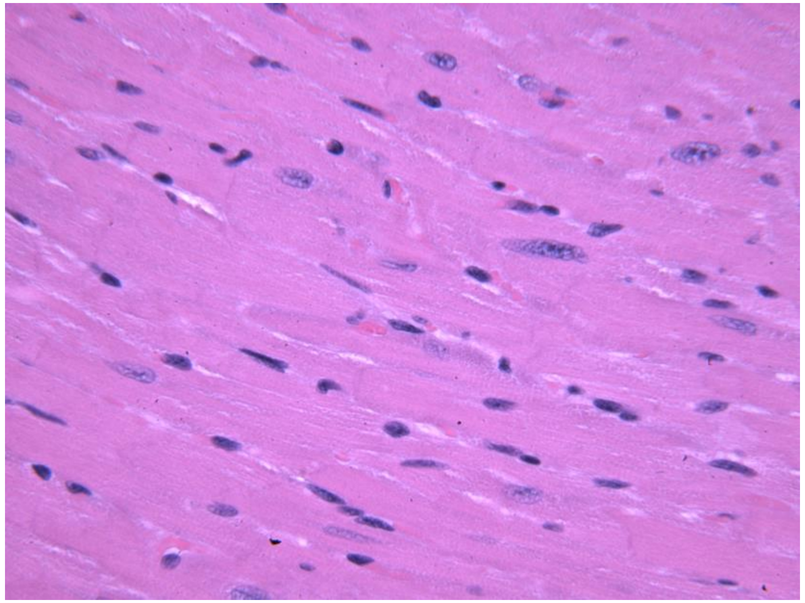

| | 10x | 40x |
|---|---|---|
| Perivascular Edema (10X) | 0 | |
| Myofibril Derangement (40X) | | 1 |
| Infiltration of Neutrophils (10X) | 1 | |
| Nuclear Hydrops (40X) | | 0 |
| Total | | |

**Scores**
**0 = no injury**
**1 = minimal (0-25%of the section)**
**2 = mild (25-50%)**
**3 = significant (50-75%)**
**4 = severe (more than 75%)**

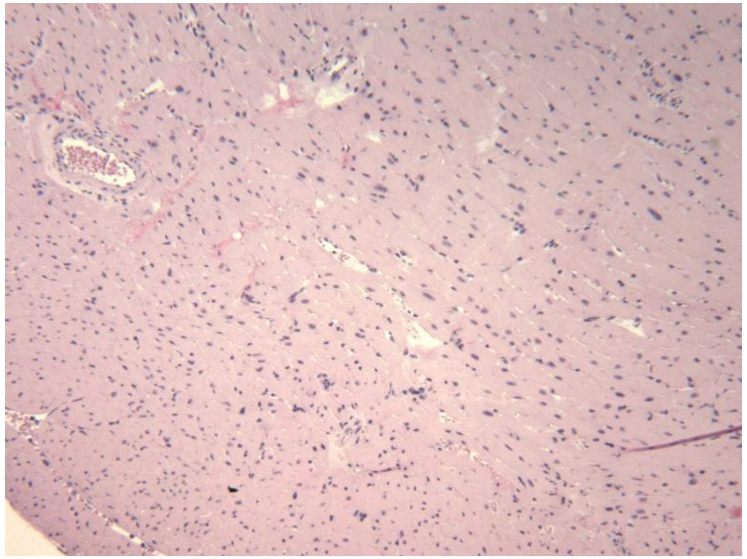

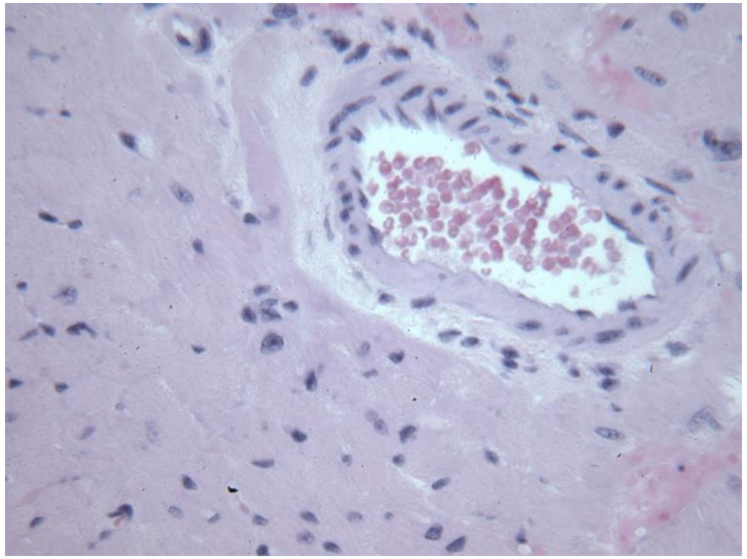

| | 10x | 40x |
|---|---|---|
| Perivascular Edema (10X) | 3 | |
| Myofibril Derangement (40X) | | 3 |
| Infiltration of Neutrophils (10X) | 3 | |
| Nuclear Hydrops (40X) | | 3 |
| Total | | |

**<u>Scores</u>**
**0 = no injury**
**1 = minimal (0-25%of the section)**
**2 = mild (25-50%)**
**3 = significant (50-75%)**
**4 = severe (more than 75%)**

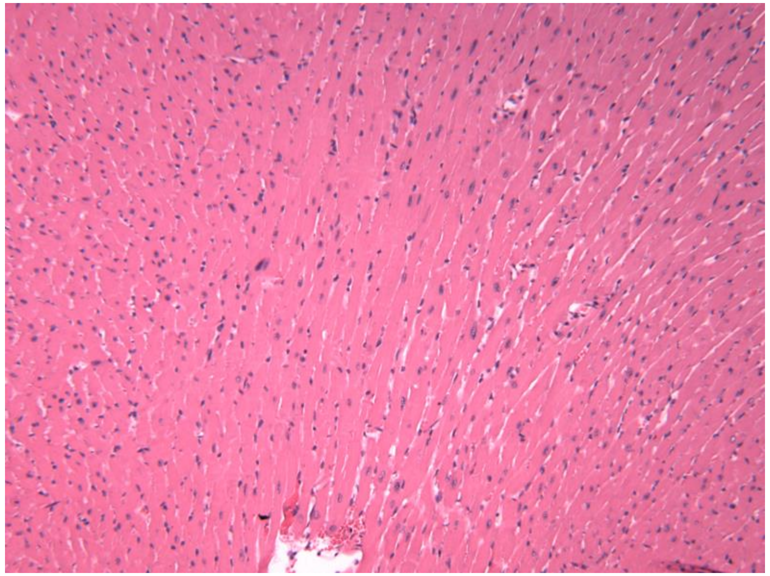

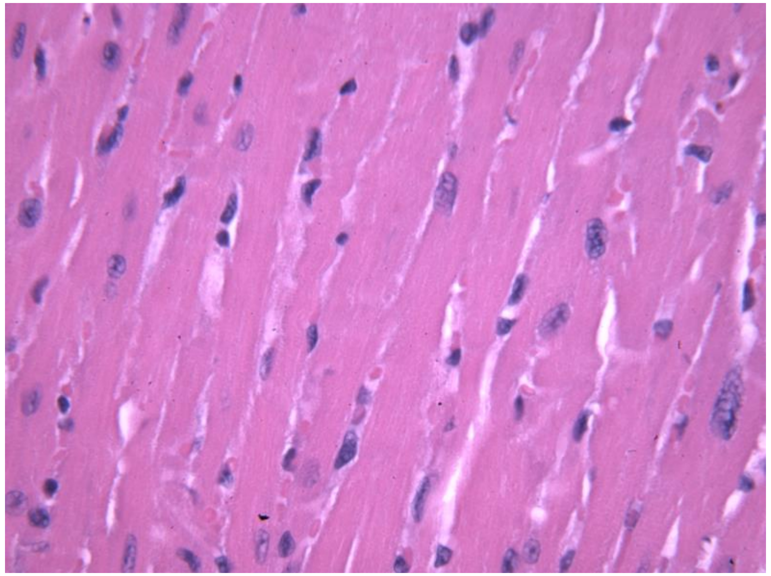

| | 10x | 40x |
|---|---|---|
| Perivascular Edema (10X) | 1 | |
| Myofibril Derangement (40X) | | 1 |
| Infiltration of Neutrophils (10X) | 2 | |
| Nuclear Hydrops (40X) | | 0 |
| Total | | |
