## Supplementary material for "KF4 anti-CELA1 Antibody and Purified α1-Antitrypsin Have Similar but Not Additive Efficacy in Preventing Emphysema in Murine α1-Antitrypsin Deficiency": Online Data Supplement-4

### Histological signs of lung injury

**Healthy Lung Tissue**

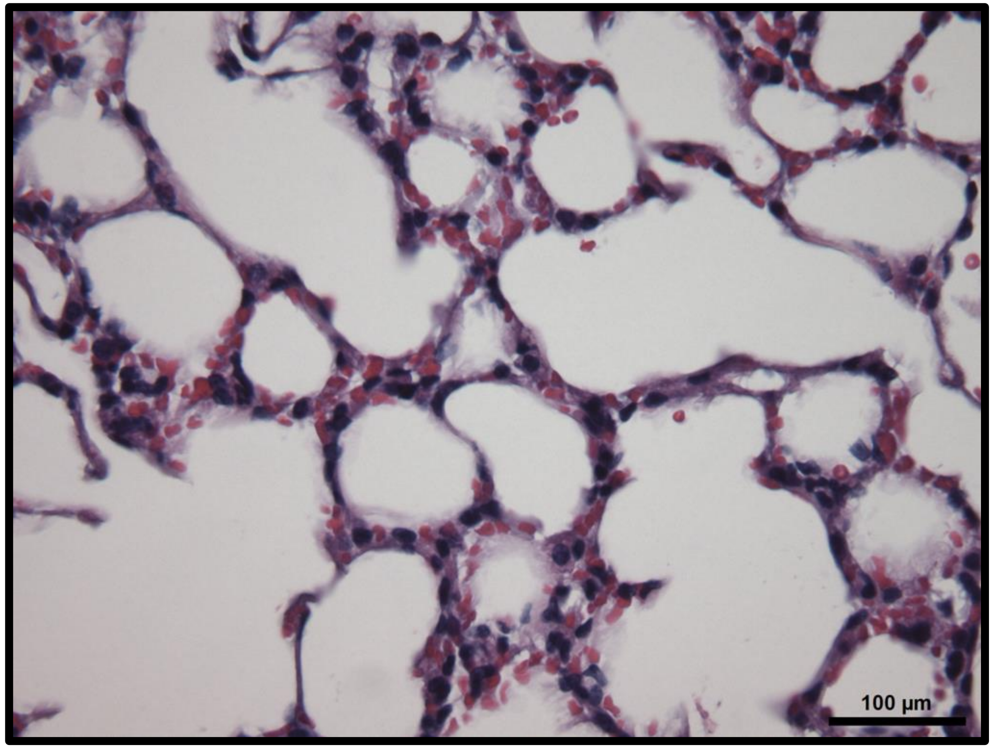

**Septic Lung Tissue**

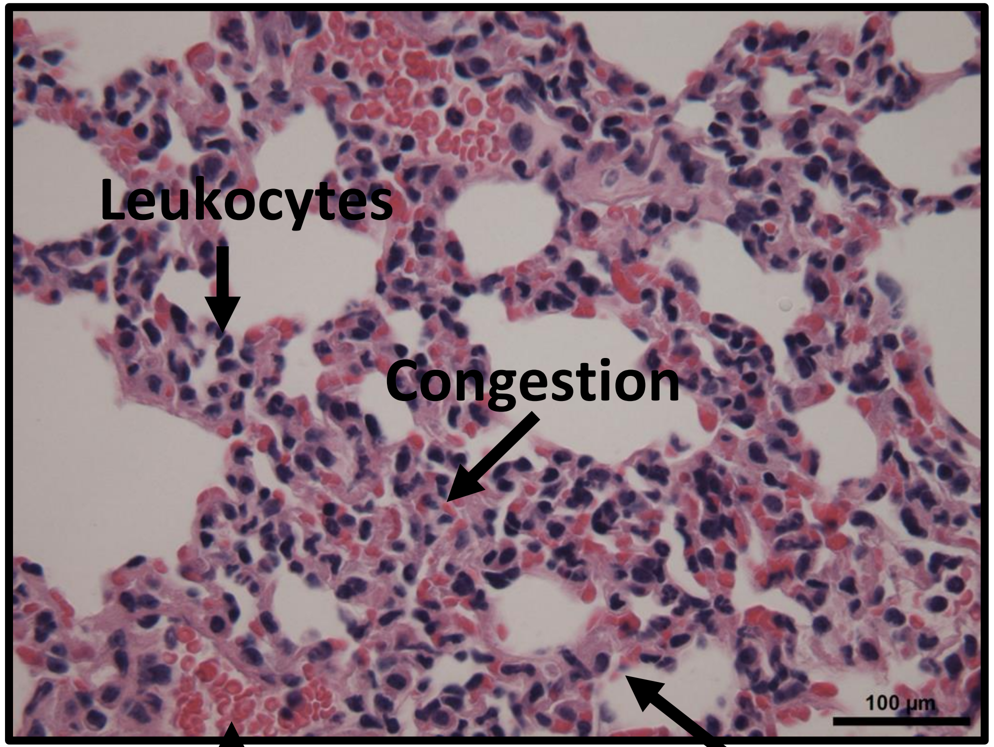

**Hemorrhage**

**Thickened Cell Walls**

**<u>Scores</u>**
**0 = no injury**
**1 = minimal (0-25%of the section)**
**2 = mild (25-50%)**
**3 = significant (50-75%)**
**4 = severe (more than 75%)**

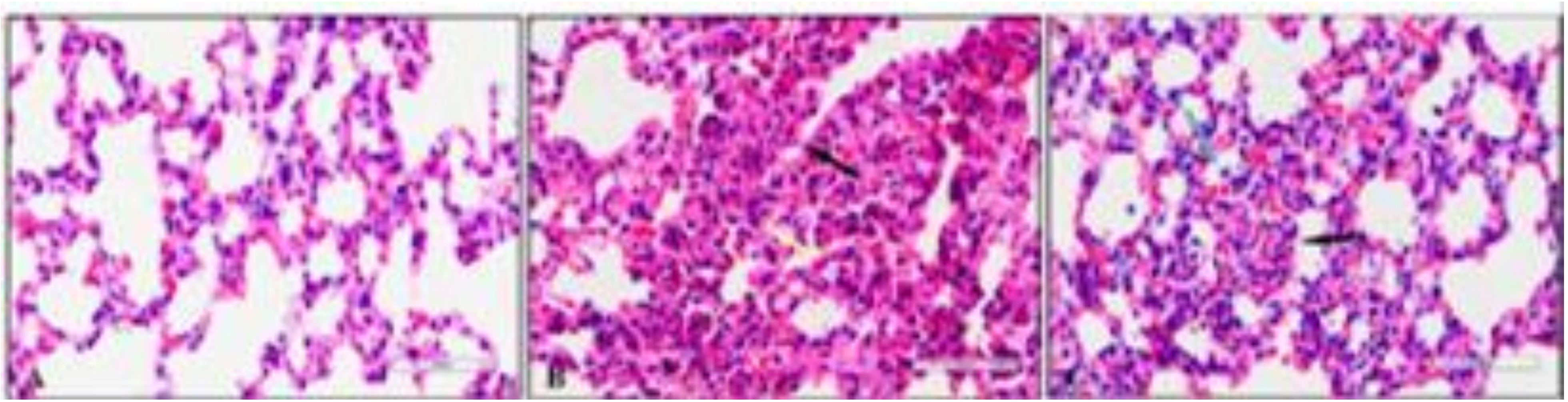

| | | | |
|---|---|---|---|
| Alveolar Congestion or Reduction of Alveolar Space | 1 | 3 | 2 |
| Hemorrhage | 1 | 3 | 3 |
| Infiltration of Leukocytes into Airspace or Alveolar Walls | 2 | 3 | 3 |
| Thickness of Alveolar Wall or Hyaline Membrane Formation | 2 | 4 | 3 |
| Total | 6 | 13 | 11 |

**<u>Scores</u>**

**0 = no injury**
**1 = minimal (0-25%of the section)**
**2 = mild (25-50%)**
**3 = significant (50-75%)**
**4 = severe (more than 75%)**

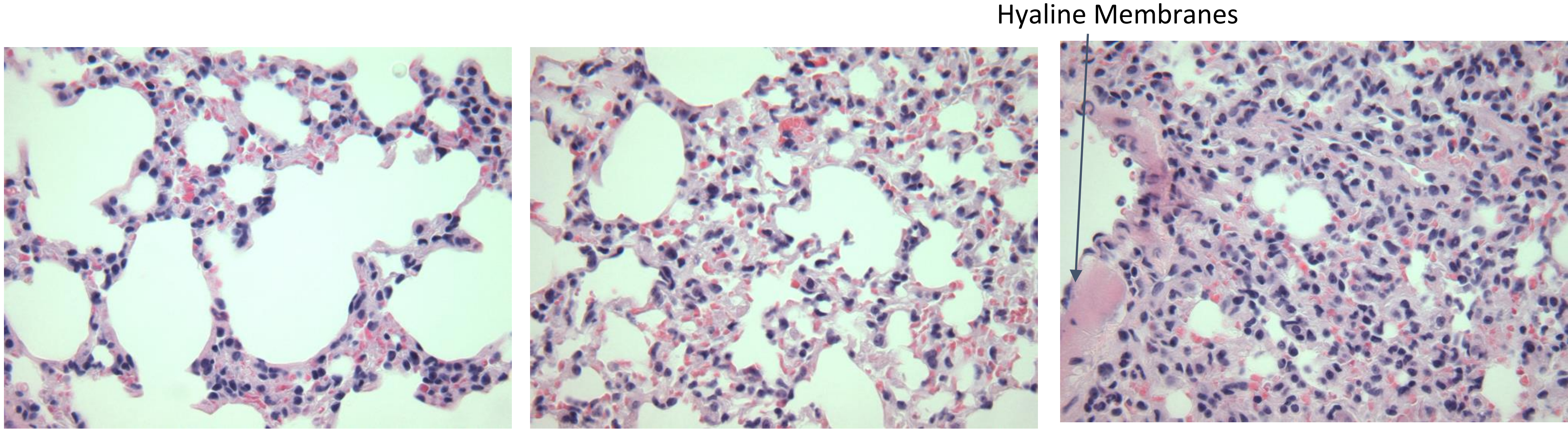

| | | | |
|---|---|---|---|
| Alveolar Congestion or Reduction of Alveolar Space | 1 | 2 | 3 |
| Hemorrhage | 0 | 1 | 1 |
| Infiltration of Leukocytes into Airspace or Alveolar Walls | 2 | 2 | 3 |
| Thickness of Alveolar Wall or Hyaline Membrane Formation | 1 | 2 | 4 |
| Total | 4 | 7 | 11 |

**<u>Scores</u>**

**0 = no injury**
**1 = minimal (0-25%of the section)**
**2 = mild (25-50%)**
**3 = significant (50-75%)**
**4 = severe (more than 75%)**

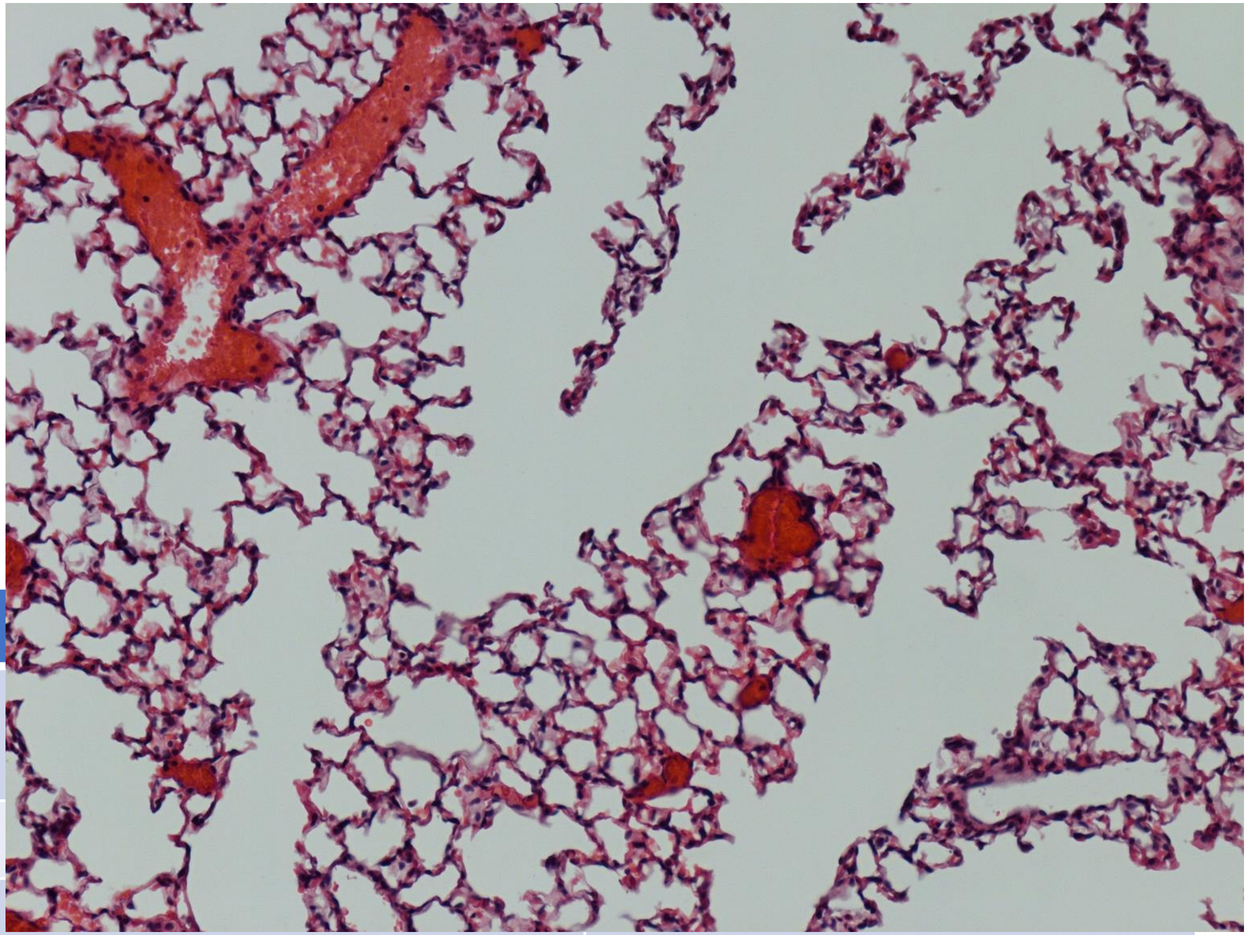

| | |
|---|---|
| Alveolar Congestion or Reduction of Alveolar Space | 0 |
| Hemorrhage | 0 |
| Infiltration of Leukocytes into Airspace or Alveolar Walls | 1 |
| Thickness of Alveolar Wall or Hyaline Membrane Formation | 2 (hyaline membranes) |
| Total | 3 |
